# Accelerated Missense Mutation Identification in Intrinsically Disordered Proteins using Deep Learning

**DOI:** 10.1101/2024.07.07.602404

**Authors:** Swarnadeep Seth, Aniket Bhattacharya

## Abstract

We use a combination of Brownian dynamics (BD) simulation results and Deep Learning (DL) strategies for rapid identification of large structural changes caused by missense mutations in intrinsically disordered proteins (IDPs). We used ∼ 6500 IDP sequences from MobiDB database of length 20 − 300 to obtain gyration radii from BD simulation on a coarse-grained single bead amino acid model (HPS2 model) used by us and others [Seth *et al*. J. Chem. Phys. **160**, 014902 (2024), Dignon *et al*. PLOS Comp. Biology, 14, 2018, Tesei *et al*. PNAS, 118, 2021] to generate the training sets for the DL algorithm. Using the gyration radii ⟨*R*_*g*_⟩ of the simulated IDPs as the training set, we develop a multilayer perceptron neural net (NN) architecture that predicts the gyration radii of 33 IDPs previously studied using BD simulation with 97% accuracy from the sequence and the corresponding parameters from the HPS model. We now utilize this NN to predict gyration radii of every permutation of missense mutations in IDPs. Our approach successfully identifies mutation-prone regions that induce significant alterations in the radius of gyration when compared to the wild-type IDP sequence. We further validate the prediction by running BD simulations on the subset of identified mutants. The neural network yields a (10^4^ − 10^6^)-fold faster computation in the search space for potentially harmful mutations. Our findings have substantial implications for rapid identification and understanding diseases related to missense mutations in IDPs and for the development of potential therapeutic interventions. The method can be extended to accurate predictions of other mutation effects in disordered proteins.

## I. Introduction

Any change in the amino acid sequence in a protein may result in non-trivial changes in its physical properties, such as, local stiffness which will affect subchain dynamics and proximity effects, its overall conformational ensemble, and dynamics that will ultimately impact its assigned biological tasks [1]. The smallest unit of that change is a missense mutation, where an amino acid in a given protein is substituted by another [2, 3]. In particular, missense mutations in intrinsically disordered proteins (IDPs) have been implicated in playing crucial roles in many diseases, such as Alzheimer’s disease (AD), Parkinson’s disease (PD), Type II Diabetes and many others [4–6]. Half of the human cancer is due to the mutation in p53. A common feature of ADs and PDs is the growth of protein droplets, resulting in liquid-liquid phase separation in IDPs [7–11], a subject which has gained considerable attention in last decade as they offer significant challenges in understanding how universal aspects of the IDPs are altered by sequence specific properties.

For an IDP of length L there are approximately L× (# of Amino Acids ∼20) ≃ 20L possible missense mutations. One of the significant challenges in the field is to narrow down and identify those missense mutations that are lethal, as these mutations are key to understanding disease mechanisms and developing targeted therapies [3]. Unlike folded proteins an increasing number of IDPs listed in the databases [12, 13] have amply demonstrated violation of the structure-function dogma as these flexible proteins sample and dynamically evolve in their vast conformational ensemble making it difficult to study experimentally. Small angle X-Ray scattering (SAXS) [14–17], high-field NMR [18, 19] and single molecule Förster resonance energy transfer (sm-FRET) [20–24] have been used to study IDPs. It is evident that traditional experimental and computational methods are impractical due to the structural flexibility of IDPs and the vast number of potential mutations [25, 26]. Thus, there is a need for a novel approach to rapidly identify lethal mutations, understand their structural implications, and develop potential therapies. Since missense mutations may adversely affect both the conformations and dynamically evolving structures of the IDPs, a practical and direct approach to develop fundamental understanding of the physical processes involved is to study not only naturally occurring but also artificial missense mutations experimentally and using computer simulation studies [26, 27]. The available experimental results have been instrumental in validating various coarse-grained models to study IDPs [28–32]. For the simplest CG models each amino acid has been lumped into a single bead [28–30]. These CG model of IDPs have been used to study single molecule properties, liquid-liquid phase separation (LLPS) in IDPs, formation of IDP-complexes [33, 43–48] and successfully extracted some key aspects of the IDPs [24]. In addition to these single-bead amino acid models those use different hydropathy scales (please refer to the next section), the two-bead representation of IDPs (SOP-IDP model) by

Baul *et al*. [31] has converged with the single-bead model and SAXS results [17]. Recently the two-bead SOP-IDP model has been extended to account for multi-domains and IDPs [32]. A major advantage of these CG models is that there is not only no cost for carrying out expensive experiments, the calculations are reasonably fast so that one can study more IDPs and available results may narrow down and direct further experiments [33]. In a previous paper, we made a detailed investigation and comparison of two CG models of IDPs (HPS1 and HPS2) [34] and calculated the mean square error (MSE) from each model. A comparison to the most recent experimental data revealed that despite use of two different hydropathy scales, both the models converge well with the experimental gyration radii values [30]. Therefore, in this article we use HPS2 model to to carry out BD simulation for a random selection of approximately 6500 IDPs from MobiDB database with disorder score ≥ 99% whose statistical properties are shown Fig. 1. Despite the fact that many properties of the Single IDPs have been investigated, we use the gyration radii from the BD simulation as the *supervised training set* for an Artificial Neural Net (ANN) to predict how missense mutation affects the physical properties of the proteins, as it is the gyration radii data that has been tabulated the most in previous studies [35, 36]. Our studies show (i) that most often missense mutations are benign but for a few the effect of the mutation is quite radical. (ii) The ANN is several order of magnitude faster to reproduce the same results with 97% accuracy showing great potential for rapidly detecting missense mutations in IDPs implicated in human disease.

**FIG. 1.**
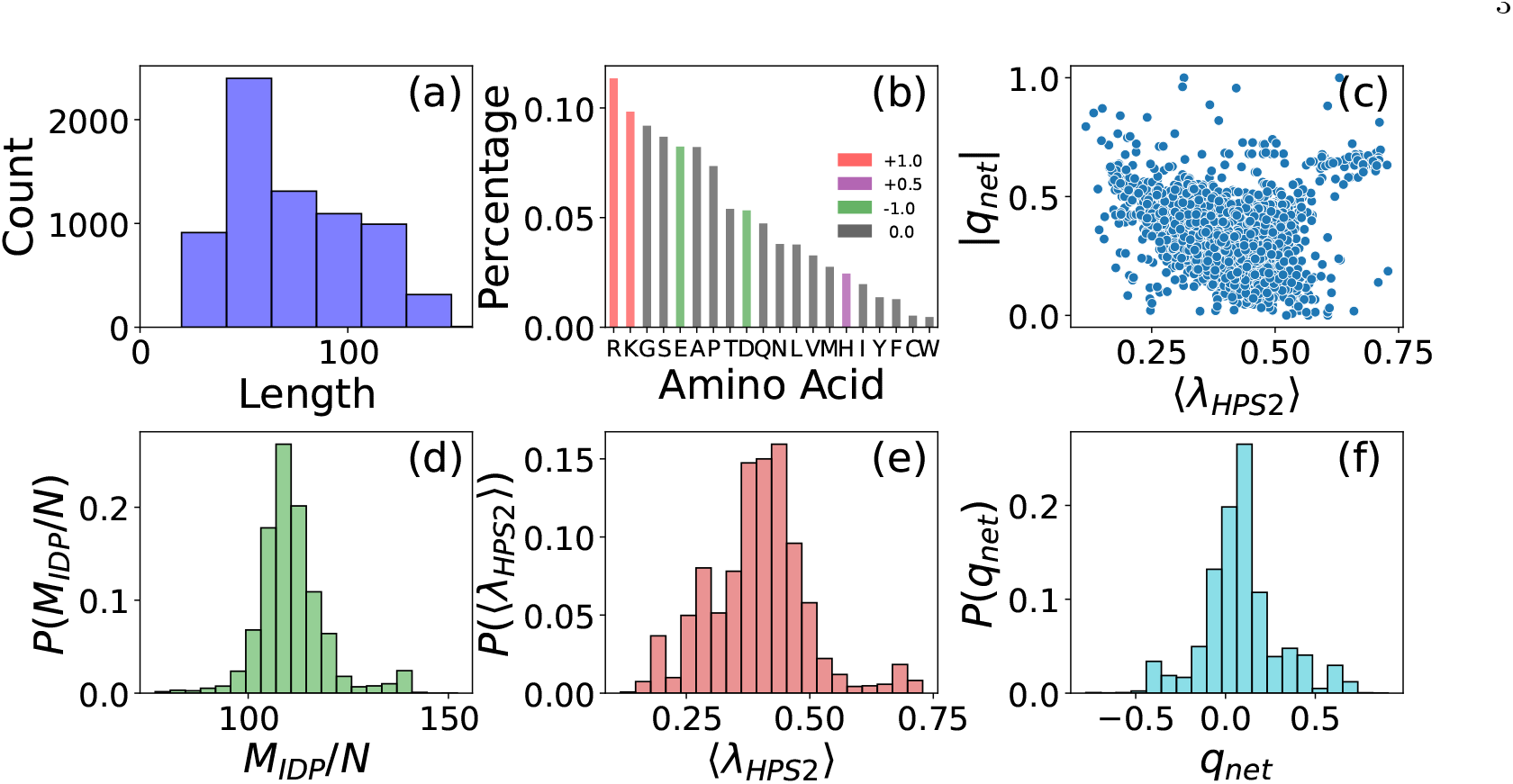
Characteristics of ∼ 6500 IDPs with ≥ 99% disorder predictor score obtained from the MobiDB [12] database. (a) Length distribution of the IDPs. (b) The distribution of the propensity of occurrence of the individual amino acids. The positive, negative, neutral amino acids are colored in red, green, and grey respectively. (c) Average hydropathy ⟨*λ*_*HPS*2_⟩ versus average net charge per residue ⟨|*q*_*net*_|⟩. (d)-(f) The distributions of mass per residue *M*_*IDP*_ */N, λ*_*HPS*2_ and *q*_*net*_ respectively.

## II. CG Model for the IDPs & ANN architecture

We provide essentials both for the HPS2 model with Ashbaugh-Hatch potential and the Neural Net architecture used to study and predict missense mutations in IDPs. Details for the HPS2 model are given in our previous paper [30].

### A. HPS2 model

In HPS2 model the amino acid residues are represented as single CG beads those interact among themselves by a modified Van der Waals interaction potential, first introduced by Ashbaugh and Hatch [37]

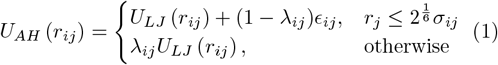

where U_*LJ*_ is the Lennard-Jones (LJ) potential,

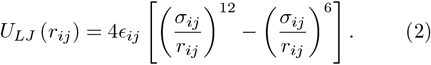

Here, 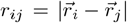 is the distance between the amino acid beads with indices i and j positioned at 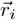 and 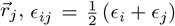 and 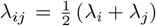 are the strength of the van der Waal interaction and average hydropathy factor between any two amino acids with indices i and j.

The hydropathy factor which is the key ingredient to differentiate interactions among IDPs. The HPS2 hydropathy scale is the same as CALVADOS introduced by Tesei *et al*. [29] using the Bayesian parameter-learning procedure to optimize the hydropathy values drawn from many different hydropathy scales [38–42] excepting the end charges are kept as they are. Here, we use HPS2 scale with optimized interaction parameter ϵ = 0.2 kcal/mol to run BD simulations to obtain the gyration radii 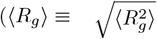 for brevity) of several thousands of IDPs to train a Neural Network architecture as discussed in the next sections. In our recent publication we demonstrated the equivalence of the HPS2 scale (with ϵ = 0.2) and the HPS1 scale (with ϵ = 0.1) due to Dignon *et al*. those produce nearly identical results with same MSE [30].

A harmonic bond potential with spring constant k_*b*_ = 8033 kJ/(mol·nm^2^) = 1920 kcal/(mol·nm^2^ (Eqn. 3)

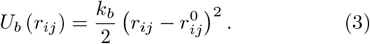

acts between two consecutive amino acid residues i and j = i ± 1. The equilibrium bond length is set to 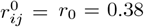 nm, the distance between α-carbon atoms for the successive amino acids. This necessitates to exclude the EV interaction among the bonded neighbors.

A screened-Coulomb (SC) interaction (Eqn. 4) acts between any two charged amino acids.

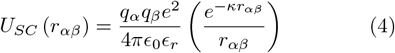

where the indices α and β refer to the subset of the indices i and j for the charged amino acids, ϵ_*r*_ is the dielectric constant of water, and κ is the inverse Debye screening length [49]. The inverse Debye length κ^−1^ is dependent on the ionic concentration (I) and expressed as

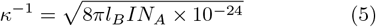

where N_*A*_ is the Avogadro’s number and l_*B*_ is the Bjerrum length,

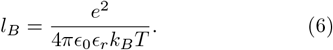

At higher temperatures, the dielectric constant typically decreases, which affects the strength of the electrostatic interactions. If the dielectric constant does not account for temperature effects, the electrostatic interactions may be overestimated, leading to unrealistic protein conformations or interactions. Hence, we implement the temperature-dependent dielectric constant of water as expressed by the empirical relation [50]

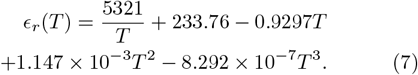

### B. Statistical properties of IDPs

We gathered approximately 6500 IDP FASTA sequences from the MobiDB database with the predicted disordered score exceeding 99%. The length of the sequences ranges between N = 20 − 150 with almost 3000 IDPs with of length between 40 to 60 as shown in Fig. 1(a). The IDPs are well known for containing a larger fraction of disorder promoting residues such as Lysine, Glutamine, Serine, Proline, Glycine, Arginine, and Glutamic acid. We gather 1400 human disordered proteome IDPs [36] and the 6500 datasets of IDPs form MobiDB dataset to calculate the propensity of occurrence of the individual amino acids and their relative probability. Fig. 1(b) confirms that positively charged Arginine and Lysine are the two most abundant residues followed by negatively charged Glutamic acid and Aspartic acid on fifth and ninth locations. We note that for large sampling of the IDPs the distributions for the mass, hydropathy and net charge per residue (Fig. 1(d)-(f)) follow a Gaussian distribution indicating pure randomness of the samples.

To conduct a detailed analysis of the statistical properties of IDPs, we carry out BD simulation for the 6500 sequences using the HPS2 hydropathy parameters to obtain the equilibrium bond length ∼ 0.382 nm so that the length of an IDP containing N Amino Acid beads L = 0.382N nm. The equilibrium radius of gyration ⟨R_*g*_⟩ has been used as the key parameter to compare with experimental results. The ⟨ R_*g*_⟩ s obtained from BD simulation on 2000 IDPs are shown in Fig. 2. A best fit produces

**FIG. 2.**
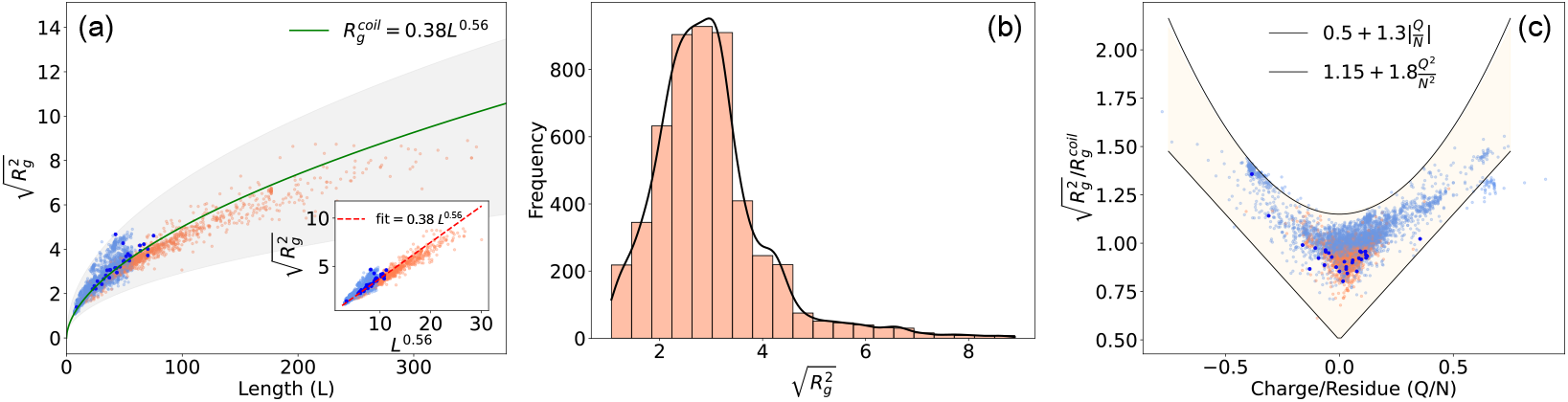
(a) Radius of gyrations of IDPs as a function of chain length. The lighter blue-shaded circles show the gyration radii of simulated IDPs using HPS2 model and the darker blue circles show the 33 IDPs from our previous study [30]. The coral-shaded circles represent 1400 human disordered proteome IDPs from [36]. The green line 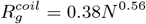 confirms the gyration radii scaling and the red dotted line in the inset plot shows gyration radii fit as a function of scaled length *L*^0.56^. (b) The distribution of 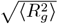 for all the ≈ 8000 IDPs. (c) The scaled gyration radii as a function of charge par residue *N/L*. The parabola and the line 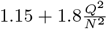 centered at 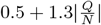 zero serve as the empirical upper and lower bounds of scaled gyration radii.

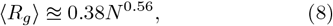

the same scaling relation that we obtained on a smaller subset of data for 33 chosen IDPs [30]. We also show two empirical bounds on the scaled gyration radii as a function of charge per residue 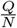 shown in Fig. 2(b). The modulus of charge/residue serves as the lower bound while the parabolic function as square of charge/residue bounds the maximum range of the scaled gyration radii. These empirical ranges will help to determine the most probable value of radius of gyration of an unknown IDPs from the net charge and number of residues.

### C. Neural Network

Artificial neural networks (ANN) are designed to emulate the human brain’s functionality by interconnecting artificial neurons. In an ANN, several layers are included in building the network architecture, with each layer consists of multiple neurons. The input layer matches the dimension of the input data, while the output layer neurons contain the information intended for learning. The intermediate layers add complexity and expressiveness to the NN, aiding in the learning process. Each neuron in every layer is connected via a weight vector to neurons in the subsequent layer. The value in each neuron is computed as a weighted sum of the values from the previous layer’s neurons, then passed through a non-linear, differentiable transformation known as the activation function. To optimize the neural network, adjustments are made to the weights assigned to all neurons, minimizing the difference between the NN’s predicted output (i.e., the output layer) and the actual output.

We train a fully connected multilayer perceptron (MLP) neural network [51] for the gyration radii prediction of disordered proteins based on a diverse set of input features based on amino acid sequence only. For the input layer of the MLP neural network we choose a subset of 23 physics-based features from a larger set those quantitatively affect and determine the gyration shape of an IDP (Fig. 3). For this supervised ANN-model training, we split the ≈ 8000 IDP dataset into 80-20 training and testing dataset. We carry out MD simulation using the HPS2 model on 6500 IDPs (and use additional ≈ 1450 gyration values from reference [36]) to obtain the radius of gyration data which serves as the true values for neural network loss calculation.

**FIG. 3.**
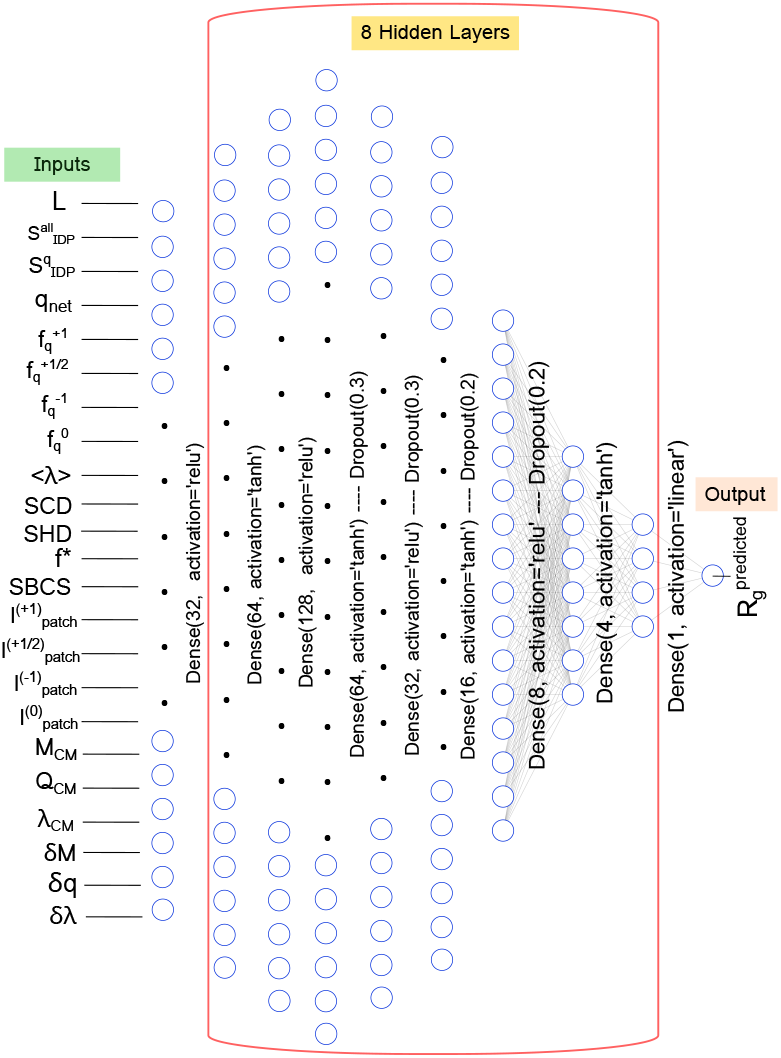
The fully connected neural network architecture to predict the gyration radii of a IDP from its sequence description. The 23 input features are obtained from the IDP FASTA sequence (please see text for the explanation of the input parameters) and are fed into an artificial neural network consists of eight hidden layers. The final layer is a single neuron that outputs the gyration radii prediction in nm.

#### 1. Physics based features as input parameters for the ANN

IDPs are mostly polyampholytes and polyelectrolytes. In addition, each amino acid bead has different mass and hydropathy indices. Therefore, it is expected the length of the chain and the distribution of the positive and negative charge resides as well as their hydropathy values will affect the conformational properties. We have used the following characteristics that depend on the sequence length, masses, charges and hydropathy indices of the amino acids making a given IDP. We define

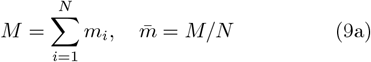

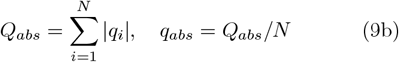

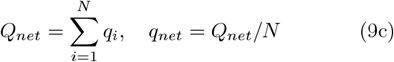

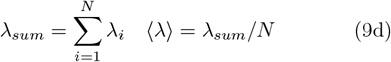

Here m_*i*_, q_*i*_, and λ_*i*_ are the mass, charge, and hydropathy index of the i^*th*^ amino acid. We include average mass per unit length 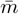 and average hydropathy ⟨λ⟩ in the training set of ANN and the quantities as itemized below.

Drawing analogy with the physical center of mass of a chain we define three quantities M_*CM*_, Q_*CM*_, λ_*CM*_, with respect to the center of the AA index (N/2) as follows:

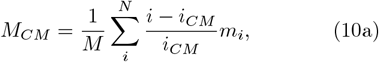

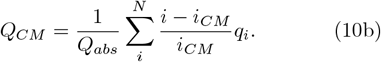

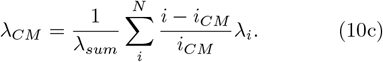

The value of i_*CM*_ is the midpoint of N, either N/2 or (N − 1)/2 depending upon the even and odd number of residues. Under this construction, the range M_*CM*_, Q_*CM*_, λ_*CM*_ can vary between (− 1, 1) that captures the asymmetry in the residue mass, charge, and hydropathy distributions regardless of the IDP length.

Likewise, we define three RMS fluctuations in mass, charge, and hydropathy

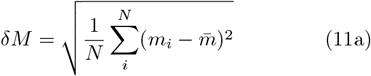

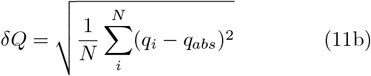

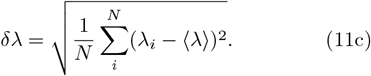

Entropy in the sequence space plays an essential role to capture the sequence dependent properties of the IDPs. We calculate the dimensionless Shannon entropy 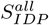 and the Shannon entropy for the charged amino acids in the sequence 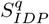 (in units of Boltzmann constant k_*B*_) of the FASTA sequence defined as follows

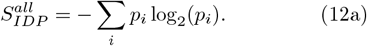

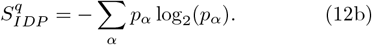

Here p_*i*_ = n_*i*_/N, where the n_*i*_ is the number of times a particular amino acid with the index i ∈ 20 occurs in the sequence. Likewise, for 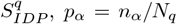, and the index α runs for the charged amino acids only with ∑*α* n_*α*_ = N_*q*_. The repetitive appearance of a particular amino acid will reduce the number of permutation and its contribution in dynamical heterogeneity will be captured by the corresponding Shannon entropy.

The other relevant quantities used to train the ANN are as follows:

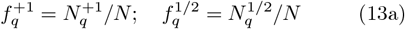

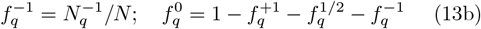

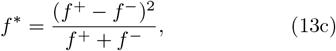

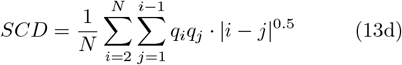

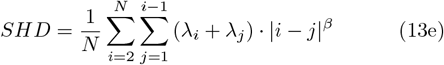

Here 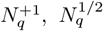, and 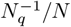 are the number of IDPs with +1, +1/2, and −1 charges, so that 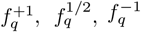 are the corresponding fractions, and 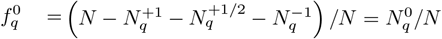 is the fraction of neutral amino acids. The quantities 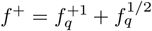 and 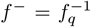 are the net positive and negative charge per residue of an IDP. The importance of charge asymmetry parameter f^*^ [43, 44] and the sequence charge decoration parameter SCD parameter (Eqn. 13d) introduced by Sawle and Ghosh [52, 53] have been discussed in the literature. In a similar spirit the Zheng *et al*. introduced hydropathy decoration parameter SHD (Eqn. 13e) [54] and provided physical arguments to show 0.95 ≤ β ≤ 1.1. They further showed that β = − 1.0 describes scaling behavior of IDPs well. We have used β = − 1 in our analysis.

The quantities

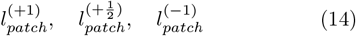

are the contiguous charge patches consisting of unit positive, half-positive, and unit negative charges respectively (please note that the IDPs have ±1 and 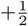 charges). 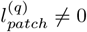, for 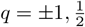 if at least two consecutive amino acids have the same charge within the charge sequence 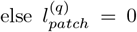. Finally, we introduce the Sequence-Based Compactness Score (SBCS) metric to evaluate the compactness of the IDP based on the distribution and clustering of high-λ residues. It is defined as

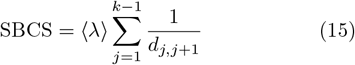

where, d_*j,j*+1_ is the is the distance between consecutive high-λ (hydrophobic) residues, indexed by j and j + 1, whose λ > 0.5 in the sequence, and k is the total number those residues. The SBCS provides insights into how high-λ residues are distributed within a sequence, and their potential role in influencing the aggregation behavior. It combines both the hydrophobicity of residues and their spatial organization to measure the overall compactness of the sequence The parameters as described in Eqns. 9(a)-9(d), 10(a)-(c), 11(a)-(c), 12(a)-(b), 13(a)-(e), 14, and 15 along with the length L of the amino acid constitute 23 important physics based features derived from the IDP sequence information as input vectors to train the MLP network as described in the next section. Since not all of them aren’t linearly independent, we have further investigated the relative weights these quantities through SHAP in Fig. 5 [55, 56].

#### 2. Neural Network Architecture

The eight hidden layers of the neural network play a crucial role in extracting intricate patterns and relationships within the input data shown in Fig. 3. The first hidden layer consists of 64 units employing the Rectified Linear Unit (ReLU) activation function, promoting non-linear transformations. Subsequently, the 2^*nd*^, 3^*rd*^, 4^*th*^·· 8^*th*^ hidden layers comprise 128, 64, 32, 16, 8 and 4 units, respectively, utilizing the Hyperbolic Tangent and ReLU activation function alternatively. This configuration is designed to facilitate the extraction of complex representations from the input features, allowing the neural network to discern relationships between the input states. The output layer of the neural network is composed of a single neuron, employing a linear activation function. This choice of activation function is tailored for regression tasks, aiming to predict a continuous output of the radius of gyration.

#### 3. Neural Network Training

The training process involves the optimization of model parameters over 1200 epochs, utilizing a batch size of 50. We employ Adam optimizer [57] with a learning and decay rate of 0.0021 and 0.0005 respectively. We jointly use Mean Squared Logarithmic Error Loss (MSLE) and Huber loss as the total loss function. The main advantage of MSLE is that it penalizes large errors more severely than Mean Squared Error (MSE) by taking the logarithm of the predicted and target values and encourages the model to predict relative errors accurately. We run the MLP network several times and incorporate dropout layers to confirm that both training and testing accuracy stay within 90 − 97% range indicating robustness of the network training within these 1200 epochs. The four dropout layers with 30%, 30%, 20%, and 20% dropout rate (shown in Fig. 3) and early stopping criteria reduce the chance of model over-fitting on the training data.

## III. Results

### A. Neural Network Predictions on Known IDPs

We validate our ANN performance by comparing the MSE error of the gyration radii values obtained from the BD simulation with those from the neural network prediction. The Table I denotes the 33 IDPs used to evaluate the model performance. The IDPs listed in Table I were studied in great detail using two HPS models and produced nearly identical results [30]. We ensured that these IDP sequences are never seen by the ANN. The gyration radii values from the BD simulation using the HPS2 model and the ANN predicted values are listed in the third and 4th column, respectively. For most of the IDPs the ANN predicts the gyration radii based on the physical characteristics of the HPS model only with an accuracy with an error ⪅ 5% with an exception of four IDPs (hNHE1cdt, synuclein, An16, and K32). We find that the use more hidden layers and the newly introduced SBSC parameter have significant effects on increasing the accuracy. The Fig. 4 shows the simulation vs. predicted radius of gyration plot for these 33 IDPs.

**TABLE 1.** Gyration Radii comparison between the simulation radius of gyration values and the neural network predicted value. The 1^*st*^ and 2^*nd*^ columns list the name and the number of amino acid residues. The simulation radius of gyration 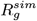 is listed in the 3^*rd*^ and neural network predicted values 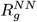 are written in the 4^*th*^ columns respectively. The last column denotes the percentage error.

| IDP Name | N | $R_g^{sim}$ (nm) | $R_g^{NN}$ (nm) | $\Delta R_g$ (%) |
| --- | --- | --- | --- | --- |
| HST5 | 24 | 1.394 | 1.468 | -5.30 |
| IN | 60 | 2.203 | 2.235 | -1.46 |
| L | 64 | 2.341 | 2.339 | 0.09 |
| CspTm | 67 | 2.223 | 2.226 | -0.13 |
| ACTR | 71 | 2.558 | 2.560 | -0.07 |
| Nucleoporin153 | 81 | 2.424 | 2.382 | 1.73 |
| SH4UD | 85 | 2.662 | 2.646 | 0.59 |
| SIC1 | 90 | 2.820 | 2.725 | 3.38 |
| p53 | 93 | 3.051 | 3.169 | -3.86 |
| CoINT | 98 | 2.715 | 2.719 | -0.14 |
| K19 | 99 | 3.065 | 2.941 | 4.05 |
| R17 | 100 | 3.075 | 3.054 | 0.69 |
| p15PAF | 111 | 3.157 | 3.130 | 0.86 |
| ProTaN | 112 | 4.671 | 4.273 | 8.52 |
| R15 | 114 | 3.145 | 3.189 | -1.40 |
| hCyp | 115 | 2.950 | 2.847 | 3.50 |
| ERMTADn | 122 | 3.210 | 3.142 | 2.11 |
| ProTaC | 129 | 4.272 | 4.273 | -0.03 |
| K18 | 130 | 3.446 | 3.362 | 2.45 |
| hNHE1cdt | 131 | 3.485 | 3.164 | 9.21 |
| sNase | 136 | 3.682 | 3.560 | 3.30 |
| synuclein | 140 | 3.787 | 3.537 | 6.61 |
| FhuA | 143 | 3.484 | 3.449 | 1.00 |
| K17 | 145 | 3.705 | 3.572 | 3.60 |
| K27 | 167 | 4.171 | 3.975 | 4.70 |
| K10 | 168 | 3.936 | 3.894 | 1.06 |
| K16 | 176 | 4.225 | 4.005 | 5.21 |
| K25 | 185 | 4.012 | 3.798 | 5.33 |
| An16 | 185 | 3.721 | 3.288 | 11.64 |
| K32 | 198 | 4.611 | 4.273 | 7.33 |
| OPN | 219 | 4.436 | 4.484 | -1.07 |

**FIG. 4.**
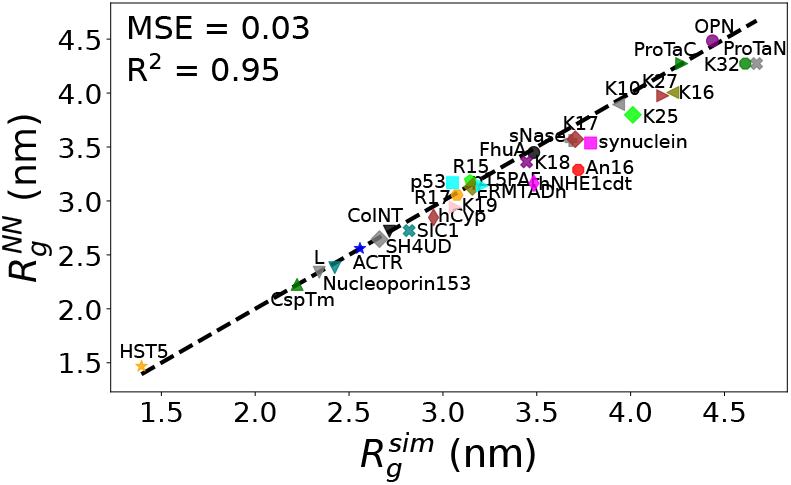
Gyration radii of each of the 33 IDPs is obtained from the BD simulation and from the neural network. The IDPs are represented with symbols in different colors. The black dashed line denotes the line of unit slope. The mean square error (MSE) is calculated from the average squared deviation of NN-predicted gyration values from the BD simulations. *R*^2^ is used for calculating the correlation coefficient.

**FIG. 5.**
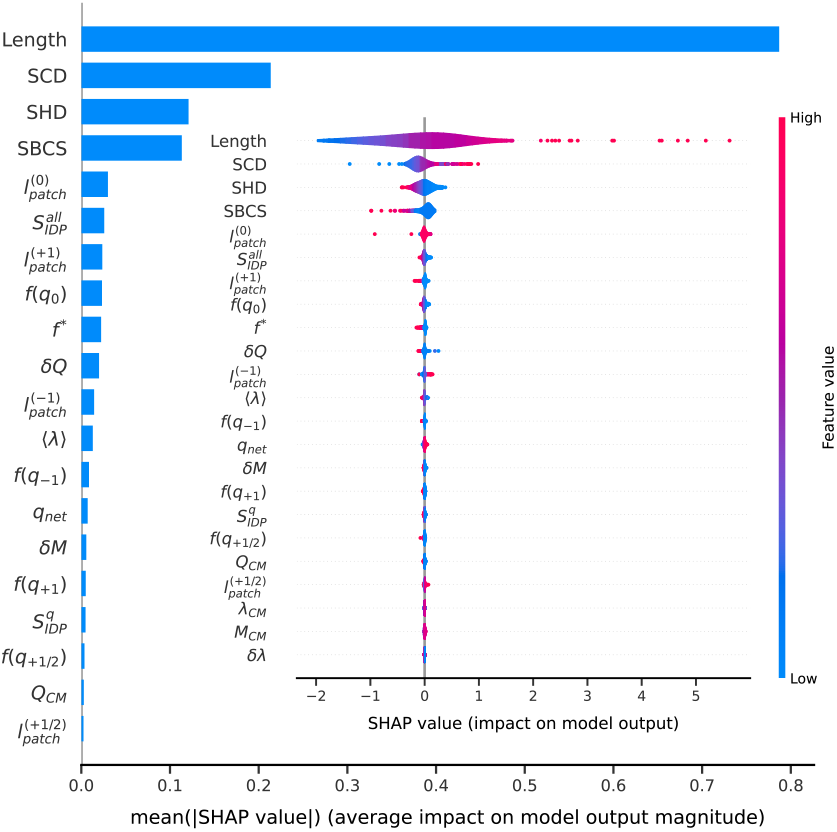
SHAP value analysis shows the feature importance in descending order. The inset (violin) plots quantitatively describe the individual feature impact on gyration radii.

### B. Identifying the feature importance using Explainable AI

Neural network models are often described as black-box models due to their use of nonlinear activation functions and multiple hidden layers, which make it almost impossible to determine how the input features contribute to meaningful outcomes, even if the model’s accuracy is high. To interpret how the model arrives at its predictions, we borrow the idea of Shapley values [55] from game theory to explain the workings of our neural network model. Shapley values (named after Lloyd Shapley) provide a method to fairly distribute the “payout” (prediction) among the features (players) based on their contributions. We utilize SHAP analysis [56] to understand our neural network’s decision-making process by quantifying the importance of each input feature and their interactions.

Fig. 5 shows the SHAP analysis of relative weights of 23 input features on the predicted gyration radii using ANN. Each point on the violin plots (inset) represents a SHAP value for one of 23 features corresponding to a single data point, with a color (red/blue for high/low values). As expected, since IDPs are described as ⟨R_*g*_ ⟩≊ 0.38L^0.56^ (Eqn. 8), SHAP predicts that length L of the IDP is the most important feature which commensurate to polymer physics principle with high values of L (indicated by red) correspond to positive SHAP values. SHAP also brings out the relative wights of various quantities pertaining to the charges on the gyration radii which is not easily interpretable or less obvious. For example, SHAP predicts that after the IDP length L, the next most important contribution comes from SCD and SHD. SCD has a positive impact on the ⟨R_*g*_⟩, with higher values increasing the ⟨R_*g*_⟩ while lower SCD slightly decreases the ⟨R_*g*_⟩. For the case of SHD this trend it opposite to that of SCD. The 4^*th*^ significant parameter we find is the SBCS introduced by us. This parameter has comparable to that of SHD with similar trend. Other features such as dQ,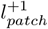, and 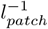 also affect the gyration radii but to a lesser extent.The important fact is that the SHAP analysis will promote further physics based research to come up with a justification a particular quantity is more/less important than another one.

### C. Identifying the change in gyration radii associated with Missense Mutations

IDPs are frequently subject to missense mutations, wherein a singular amino acid is substituted by one of the remaining 19 amino acids. Pathogenic missense variants disrupt protein function and reduce organismal fitness, while benign missense variants have limited effects. Consequently, a comprehensive exploration of all potential mutant sequences for a given IDP is imperative. Traditional Molecular Dynamics (MD) simulations for calculating gyration radii across the entire spectrum of mutant sequences are inherently time-consuming. To circumvent this computational bottleneck, we employ our trained ANN model, which significantly expedites calculation of gyration radii for all conceivable mutants. This innovative approach enables a significant reduction in computational time, rendering our methodology highly efficient for exhaustive mutational analyses within the realm of IDPs. It is worth noting that a large variation in ⟨R_*g*_⟩ may not necessarily imply a pathogenic variant and worthy of further studies.

Finally, we to demonstrate the eficacy of the ANN prediction we make a one-to-one comparison of the 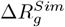 with 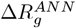 in a color map for the HST5, IDP of 24 amino acid residues shown side by side in Fig. 6 and note that the corresponding overlap integral for each missense mutation is as large as 80%. This opens the scope that the neural network performance can be improved by training with a high amount of simulation/experimental data if available along with new sequence based functions, such as SBSC introduced in this paper.

**FIG. 6.**
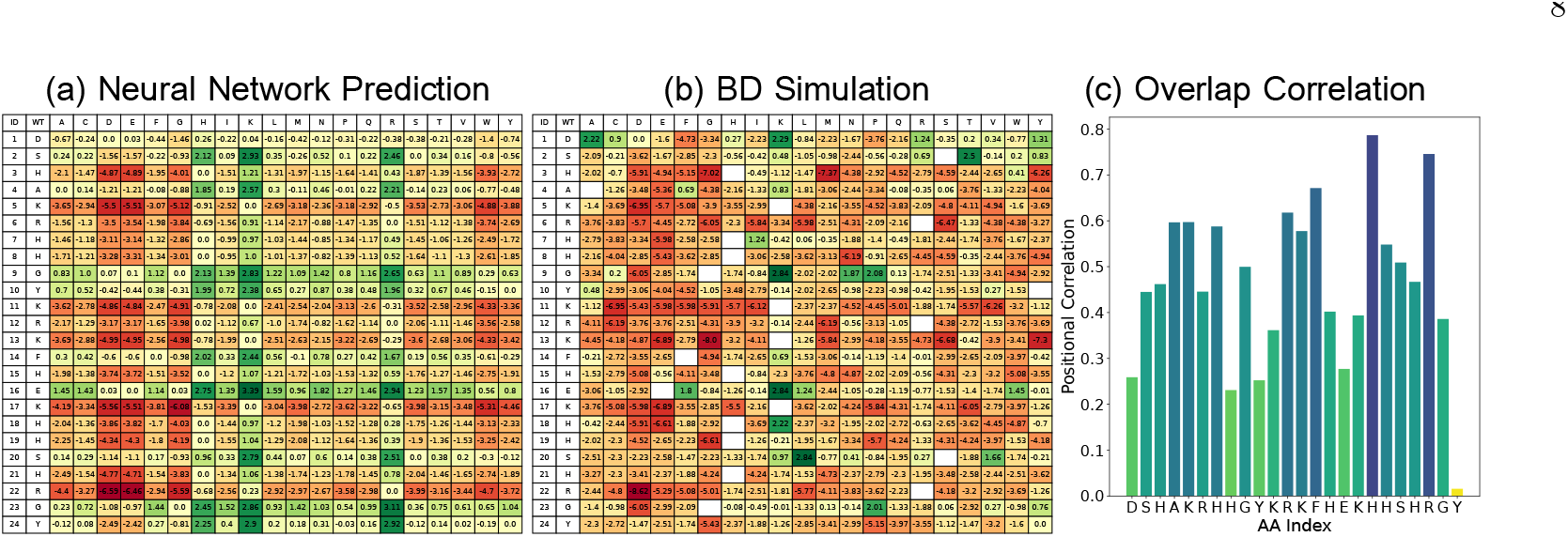
(a) The checker-board heat-map of the percentage change in the gyration radii of HST5 with the point mutation from neural network prediction. The first column of the map denotes the wild type sequence and the point mutations are listed in the first row. The corresponding percentage change in the gyration radii is shown in red to green colors. The dark red denotes the mutant gyration radii is greater than the wild-type sequence and vice versa for the green boxes. (b) denotes the same changes obtained from the BD simulation. (c) compares the correlation between the neural network prediction and the BD simulations results.

### D. Summary and Outlook

To summarize, we have provided a straight forward, reasonably accurate and computationally efficient algorithm to identify missense mutations in IDPs using a combination of simulation results from a well tested CG model of IDPs and feeding this information to a ANN. We have shown that the ANN predicts the same with 97% accuracy and six orders of magnitude faster than the BD algorithm. The algorithm is open for significant and continuous improvement as (i) more and more data from the BD simulation data backed up by experiments become available to train the ANN and (ii) choice of new parameters for the training sets. The heat map of the results shows that a large fraction of the mutations are less significant while only a few undergo a large variation in sizes. Thus, the use of ANN to rapidly identify this small subset is a key and important element in our findings. This will allow to go back to these few mutated species and carry out BD simulation to gain a fundamental understanding of the accompanying changes not attainable by machine learning algorithm only.

Though, the neural network traditionally viewed as a black-box, we utilize SHAP analysis to understand the degree of impact of individual physics based feature on gyration radii prediction and in turns sheds lights on the inner workings of our neural network model. We showed that the length of IDPs is the most significant predictor of the radius of gyration, followed by the SCD, SHD, and the SBSC parameter introduced in this paper. This transparency not only aligns with polymer physics principles but also enhances the interpretability and trustworthiness of our NN model. This highlight the importance of explainable AI in understanding and validating complex neural network models with more confidence.

We want to conclude with some remark making connection to this work with other work where Physics based model along with ML algorithms have been used to explore IDPs [35, 36, 58–61]. It is immensely practical to increase the accuracy of predictions the conformational properties of the IDPs as much as possible directly from the sequence itself, although other physical features will be needed for a more comprehensive physical understanding. From the ML perspective there can be multiple approaches in terms of different deep learning strategies, choice of training sets, use of simulation and experimental data, *etc*.. As an example, Chao *et al*. [60] used various regression models, *e*.*g*., L2-Linear regularized regression (LRR), Kernel ridge regression (KRR), Gaussian process regression (GPR), and free forward neural network (FNN) to compare MSEs from different regression methods. LRR, KRR, and GPR produce MSE in the range 0.93-0.94, comparable to but slightly lower than ours reported here. Janson *et al*. [61] used the well established Generative Adversarial Network (GAN) [62] to generate conformational ensembles of IDPs which they call idpGAN. idpGAN can generate physically realistic conformational ensembles of proteins without encountering kinetic barriers in an actual MD simulation and therefore, it is extremely fast and complements ours and other ML methods used by Chao *et al*. A combination of different ML algorithms along with the data generated from simulation of different level of granurality will be a practical approach to speed up the investigations [58]. Lotthammer *et al*. introduced a deep learning package called ALBATROSS [35] using a bidirectional recurrent neural network with long short-term memory cells (LSTM-BRNN) to study a total of 41,202 IDPs and IDRs (19,075 naturally occurring IDRs and 22,127 rationally designed IDRs). However, Lotthammer *et al*. used ABSINTH [63] but not the HPS2 and therefore, a comparison of the results will reinforce the findings to the IDP community. In a related study Tesei *et al*. used CALVADOS and Machine learning model to study 28,058 occurring in human proteome to predict apparent scaling exponents [36]. Tesei *et al*. found only a small shift in the histogram of the entropy in benign and pathogenic variants. By calculating the effective Flory exponents for proteins from Clinvar database [64] they found compactness of benign and pathogenic variants are statistically insignificant. Google DeepMind recently reported prediction of pathogenic mutations in α-synuclein [3]. The final prediction of α-Missense is a pathogenicity score, which reflect the likelihoods of mutations to cause disease, rather than predicted changes in protein structures [65]. While the algorithm may well be very efficient, this provides a probability score without providing any physical justification. Our approach (an similar approaches recently used by others) on the contrary based on physical models of IDPs which have been used to disease progression and liquid-liquid phase separation in living systems with the parameters validated by experiments. Admittedly, predictions solely based on sequence-gazing and ML algorithms have their limitations, will not be able to capture the physical scenario which will require further knowledge of the dynamics. For example, visualization of free energy landscape has revealed important information about fibril formation in Amyloid-beta [66, 67]. However, collective explainable ML methods without carrying out expensive simulation with the SHAP analysis is extremely useful in narrowing down the key parameters for vast number of IDPs in different physical environment for time consuming simulation studies. Our studies of missense mutations is a practical step in that direction which we believe will promote further work for broad general application in rapidly identifying harmful mutations in proteins and ultimately clinical variant effect predictions.

## Supporting information

Mutation heat-map CspTM

Mutation heat-map IN

Mutation heat-map p53

Mutation heat-map ProTaC

Mutation heat-map ProTaN

Mutation heat-map Synuclein

## Acknowledgments

The research at UCF has been supported by a DARPA AI-BTO (HR00112530047) award. All computations were carried out using STOKES High-Performance Computing Cluster at UCF. We thank both the anonymous reviewers for a thorough reading of the original manuscript and comments.

## Notes

### Competing Interest Statement

The authors have declared no competing interest.

### Summary of Updates

The manuscript has been updated to include revised graphs with more data point from new simulation runs and inclusion of additional references.

