## Supplementary material for "Accelerated Missense Mutation Identification in Intrinsically Disordered Proteins using Deep Learning": Mutation heat-map p53

| ID | WT | A | C | D | E | F | G | H | I | K | L | M | N | P | Q | R | S | T | V | W | Y |
|---|---|---|---|---|---|---|---|---|---|---|---|---|---|---|---|---|---|---|---|---|---|
| 1 | M | 0.03 | 0.02 | 0.08 | 0.06 | -0.08 | -0.17 | -0.07 | 0.04 | 0.16 | 0.11 | 0.0 | 0.08 | 0.12 | 0.05 | -0.04 | 0.07 | 0.08 | 0.08 | -0.45 | -0.26 |
| 2 | E | -1.07 | -1.5 | 0.04 | 0.0 | -1.96 | -2.03 | -2.72 | -1.78 | -3.01 | -1.22 | -1.93 | -1.2 | -1.26 | -1.39 | -3.64 | -1.36 | -1.25 | -1.36 | -2.31 | -2.31 |
| 3 | E | -1.03 | -1.46 | 0.04 | 0.0 | -1.86 | -1.93 | -2.66 | -1.65 | -2.94 | -1.18 | -1.82 | -1.16 | -1.22 | -1.35 | -3.57 | -1.33 | -1.21 | -1.32 | -2.27 | -2.23 |
| 4 | P | 0.02 | -0.08 | 1.07 | 1.02 | -0.28 | -0.34 | -0.61 | -0.13 | -0.74 | 0.02 | -0.23 | 0.04 | 0.0 | -0.07 | -1.33 | -0.05 | 0.02 | -0.04 | -0.62 | -0.47 |
| 5 | Q | 0.11 | -0.01 | 1.13 | 1.07 | -0.22 | -0.28 | -0.53 | -0.06 | -0.66 | 0.11 | -0.17 | 0.12 | 0.08 | 0.0 | -1.2 | 0.02 | 0.1 | 0.04 | -0.54 | -0.41 |
| 6 | S | 0.1 | -0.02 | 1.09 | 1.03 | -0.24 | -0.3 | -0.53 | -0.08 | -0.64 | 0.09 | -0.19 | 0.11 | 0.06 | -0.01 | -1.18 | 0.0 | 0.08 | 0.03 | -0.55 | -0.43 |
| 7 | D | -1.02 | -1.5 | 0.0 | -0.04 | -1.96 | -2.02 | -2.7 | -1.69 | -2.87 | -1.2 | -1.94 | -1.15 | -1.27 | -1.4 | -3.6 | -1.38 | -1.22 | -1.36 | -2.41 | -2.41 |
| 8 | P | 0.05 | -0.08 | 0.99 | 0.93 | -0.3 | -0.36 | -0.59 | -0.14 | -0.68 | 0.03 | -0.25 | 0.05 | 0.0 | -0.07 | -1.26 | -0.05 | 0.02 | -0.03 | -0.64 | -0.5 |
| 9 | S | 0.11 | -0.02 | 1.09 | 1.03 | -0.25 | -0.31 | -0.52 | -0.09 | -0.62 | 0.1 | -0.2 | 0.12 | 0.06 | -0.01 | -1.16 | 0.0 | 0.09 | 0.03 | -0.56 | -0.44 |
| 10 | V | 0.1 | -0.04 | 1.04 | 0.98 | -0.27 | -0.32 | -0.53 | -0.1 | -0.6 | 0.08 | -0.22 | 0.1 | 0.04 | -0.02 | -1.15 | -0.01 | 0.07 | 0.0 | -0.6 | -0.46 |
| 11 | E | -0.91 | -1.42 | 0.04 | 0.0 | -1.9 | -1.95 | -2.57 | -1.62 | -2.68 | -1.12 | -1.89 | -1.06 | -1.19 | -1.32 | -3.48 | -1.3 | -1.13 | -1.28 | -2.33 | -2.35 |
| 12 | P | 0.06 | -0.08 | 0.98 | 0.92 | -0.31 | -0.36 | -0.58 | -0.14 | -0.63 | 0.04 | -0.26 | 0.06 | 0.0 | -0.07 | -1.22 | -0.06 | 0.03 | -0.03 | -0.66 | -0.51 |
| 13 | P | 0.06 | -0.08 | 0.99 | 0.93 | -0.31 | -0.36 | -0.58 | -0.15 | -0.64 | 0.04 | -0.26 | 0.06 | 0.0 | -0.07 | -1.24 | -0.06 | 0.03 | -0.03 | -0.66 | -0.52 |
| 14 | L | 0.03 | -0.11 | 0.94 | 0.88 | -0.34 | -0.39 | -0.62 | -0.18 | -0.68 | 0.0 | -0.29 | 0.03 | -0.03 | -0.1 | -1.31 | -0.08 | -0.0 | -0.06 | -0.71 | -0.56 |
| 15 | S | 0.13 | -0.02 | 1.07 | 1.01 | -0.26 | -0.31 | -0.5 | -0.1 | -0.56 | 0.1 | -0.22 | 0.13 | 0.06 | -0.01 | -1.12 | 0.0 | 0.1 | 0.03 | -0.57 | -0.45 |
| 16 | Q | 0.15 | -0.0 | 1.08 | 1.01 | -0.25 | -0.3 | -0.48 | -0.08 | -0.52 | 0.12 | -0.2 | 0.14 | 0.08 | 0.0 | -1.08 | 0.02 | 0.11 | 0.05 | -0.56 | -0.44 |
| 17 | E | -0.85 | -1.37 | 0.04 | 0.0 | -1.92 | -1.95 | -2.48 | -1.65 | -2.52 | -1.07 | -1.92 | -0.99 | -1.14 | -1.27 | -3.39 | -1.25 | -1.08 | -1.23 | -2.29 | -2.35 |
| 18 | T | 0.04 | -0.1 | 0.92 | 0.86 | -0.35 | -0.4 | -0.59 | -0.19 | -0.62 | 0.02 | -0.32 | 0.04 | -0.02 | -0.09 | -1.33 | -0.08 | 0.0 | -0.05 | -0.69 | -0.6 |
| 19 | F | 0.47 | 0.29 | 1.52 | 1.45 | 0.0 | -0.04 | -0.17 | 0.2 | -0.17 | 0.43 | 0.07 | 0.46 | 0.39 | 0.31 | -0.66 | 0.32 | 0.42 | 0.35 | -0.32 | -0.2 |
| 20 | S | 0.13 | -0.02 | 1.04 | 0.98 | -0.27 | -0.32 | -0.49 | -0.11 | -0.51 | 0.1 | -0.24 | 0.13 | 0.06 | -0.01 | -1.16 | 0.0 | 0.1 | 0.03 | -0.59 | -0.5 |
| 21 | D | -0.88 | -1.39 | 0.0 | -0.04 | -2.0 | -2.03 | -2.51 | -1.72 | -2.53 | -1.09 | -2.01 | -1.01 | -1.17 | -1.31 | -3.42 | -1.28 | -1.1 | -1.26 | -2.36 | -2.42 |
| 22 | L | 0.04 | -0.11 | 0.91 | 0.85 | -0.38 | -0.43 | -0.6 | -0.2 | -0.62 | 0.0 | -0.35 | 0.04 | -0.03 | -0.1 | -1.37 | -0.09 | 0.0 | -0.06 | -0.7 | -0.62 |
| 23 | W | 0.84 | 0.54 | 2.05 | 1.99 | 0.28 | 0.26 | 0.05 | 0.43 | -0.18 | 0.71 | 0.29 | 0.75 | 0.67 | 0.58 | -0.77 | 0.6 | 0.7 | 0.62 | 0.0 | 0.08 |
| 24 | K | 0.7 | 0.52 | 1.85 | 1.77 | 0.22 | 0.16 | 0.02 | 0.43 | 0.0 | 0.67 | 0.27 | 0.71 | 0.62 | 0.53 | -0.36 | 0.55 | 0.66 | 0.58 | -0.15 | -0.02 |
| 25 | L | 0.04 | -0.11 | 0.94 | 0.88 | -0.35 | -0.4 | -0.62 | -0.19 | -1.02 | 0.0 | -0.31 | 0.04 | -0.03 | -0.1 | -1.74 | -0.09 | 0.0 | -0.06 | -0.71 | -0.6 |
| 26 | L | 0.04 | -0.11 | 0.94 | 0.86 | -0.35 | -0.4 | -0.6 | -0.19 | -0.62 | 0.0 | -0.31 | 0.04 | -0.03 | -0.1 | -1.28 | -0.09 | 0.0 | -0.06 | -0.72 | -0.59 |
| 27 | P | 0.07 | -0.08 | 0.94 | 0.88 | -0.32 | -0.37 | -0.55 | -0.16 | -0.55 | 0.04 | -0.28 | 0.07 | 0.0 | -0.07 | -1.15 | -0.06 | 0.04 | -0.03 | -0.67 | -0.55 |
| 28 | E | -0.8 | -1.3 | 0.04 | 0.0 | -1.85 | -1.88 | -2.37 | -1.55 | -2.36 | -1.0 | -1.86 | -0.92 | -1.08 | -1.22 | -3.27 | -1.19 | -1.01 | -1.16 | -2.25 | -2.31 |
| 29 | N | 0.02 | -0.13 | 0.85 | 0.79 | -0.37 | -0.42 | -0.62 | -0.21 | -0.62 | -0.02 | -0.33 | 0.0 | -0.05 | -0.12 | -1.25 | -0.11 | -0.02 | -0.08 | -0.76 | -0.63 |
| 30 | N | 0.02 | -0.13 | 0.86 | 0.8 | -0.37 | -0.42 | -0.62 | -0.21 | -0.63 | -0.02 | -0.33 | 0.0 | -0.05 | -0.12 | -1.27 | -0.11 | -0.02 | -0.08 | -0.76 | -0.63 |
| 31 | V | 0.12 | -0.04 | 1.0 | 0.93 | -0.29 | -0.34 | -0.5 | -0.12 | -0.5 | 0.08 | -0.25 | 0.11 | 0.04 | -0.03 | -1.06 | -0.02 | 0.08 | 0.0 | -0.62 | -0.5 |
| 32 | L | 0.04 | -0.11 | 0.89 | 0.83 | -0.35 | -0.4 | -0.59 | -0.19 | -0.59 | 0.0 | -0.31 | 0.04 | -0.03 | -0.1 | -1.21 | -0.09 | 0.0 | -0.06 | -0.73 | -0.59 |
| 33 | S | 0.14 | -0.02 | 1.02 | 0.96 | -0.27 | -0.32 | -0.47 | -0.1 | -0.46 | 0.1 | -0.23 | 0.14 | 0.06 | -0.01 | -1.02 | 0.0 | 0.1 | 0.03 | -0.58 | -0.48 |
| 34 | P | 0.08 | -0.08 | 0.93 | 0.87 | -0.32 | -0.37 | -0.54 | -0.16 | -0.53 | 0.04 | -0.28 | 0.08 | 0.0 | -0.07 | -1.12 | -0.06 | 0.04 | -0.03 | -0.68 | -0.55 |
| 35 | L | 0.04 | -0.11 | 0.87 | 0.81 | -0.36 | -0.4 | -0.58 | -0.19 | -0.57 | 0.0 | -0.32 | 0.04 | -0.03 | -0.1 | -1.19 | -0.09 | 0.0 | -0.06 | -0.73 | -0.6 |
| 36 | P | 0.08 | -0.08 | 0.91 | 0.86 | -0.32 | -0.37 | -0.53 | -0.16 | -0.51 | 0.04 | -0.28 | 0.08 | 0.0 | -0.07 | -1.1 | -0.06 | 0.04 | -0.03 | -0.68 | -0.55 |
| 37 | S | 0.14 | -0.02 | 0.99 | 0.93 | -0.27 | -0.32 | -0.45 | -0.1 | -0.43 | 0.1 | -0.23 | 0.14 | 0.06 | -0.01 | -0.99 | 0.0 | 0.1 | 0.03 | -0.58 | -0.49 |
| 38 | Q | 0.16 | 0.0 | 1.01 | 0.94 | -0.26 | -0.31 | -0.43 | -0.09 | -0.4 | 0.12 | -0.22 | 0.16 | 0.08 | 0.0 | -0.97 | 0.02 | 0.12 | 0.05 | -0.58 | -0.48 |
| 39 | A | 0.0 | -0.16 | 0.79 | 0.74 | -0.51 | -0.54 | -0.66 | -0.33 | -0.63 | -0.04 | -0.5 | -0.01 | -0.08 | -0.14 | -1.47 | -0.14 | -0.05 | -0.11 | -0.77 | -0.74 |
| 40 | M | 0.41 | 0.25 | 1.38 | 1.33 | -0.04 | -0.1 | -0.17 | 0.16 | -0.11 | 0.37 | 0.0 | 0.41 | 0.33 | 0.25 | -0.56 | 0.26 | 0.37 | 0.3 | -0.39 | -0.26 |
| 41 | D | -0.72 | -1.11 | 0.0 | -0.04 | -1.79 | -1.8 | -2.07 | -1.54 | -1.94 | -0.83 | -1.81 | -0.77 | -0.92 | -1.05 | -2.91 | -1.02 | -0.83 | -0.99 | -2.13 | -2.21 |
| 42 | D | -0.71 | -1.1 | 0.0 | -0.04 | -1.73 | -1.75 | -2.05 | -1.47 | -1.91 | -0.82 | -1.74 | -0.77 | -0.91 | -1.03 | -2.88 | -1.01 | -0.82 | -0.98 | -2.09 | -2.16 |
| 43 | L | 0.04 | -0.11 | 0.83 | 0.79 | -0.4 | -0.44 | -0.54 | -0.21 | -0.47 | 0.0 | -0.38 | 0.04 | -0.03 | -0.1 | -1.14 | -0.09 | 0.0 | -0.06 | -0.72 | -0.65 |
| 44 | M | 0.41 | 0.25 | 1.29 | 1.22 | -0.04 | -0.1 | -0.17 | 0.16 | -0.11 | 0.37 | 0.0 | 0.41 | 0.33 | 0.25 | -0.55 | 0.26 | 0.37 | 0.3 | -0.39 | -0.26 |
| 45 | L | 0.04 | -0.11 | 0.8 | 0.74 | -0.4 | -0.44 | -0.54 | -0.21 | -0.49 | 0.0 | -0.37 | 0.04 | -0.03 | -0.1 | -1.16 | -0.09 | 0.0 | -0.06 | -0.72 | -0.65 |
| 46 | S | 0.14 | -0.02 | 0.93 | 0.87 | -0.28 | -0.32 | -0.42 | -0.11 | -0.36 | 0.1 | -0.24 | 0.14 | 0.06 | -0.01 | -0.92 | 0.0 | 0.1 | 0.03 | -0.59 | -0.5 |
| 47 | P | 0.08 | -0.08 | 0.87 | 0.82 | -0.33 | -0.38 | -0.49 | -0.16 | -0.41 | 0.04 | -0.29 | 0.08 | 0.0 | -0.07 | -0.98 | -0.06 | 0.04 | -0.03 | -0.67 | -0.57 |
| 48 | D | -0.69 | -1.06 | 0.0 | -0.04 | -1.66 | -1.68 | -1.98 | -1.4 | -1.82 | -0.8 | -1.68 | -0.74 | -0.87 | -1.0 | -2.79 | -0.97 | -0.8 | -0.94 | -2.04 | -2.1 |
| 49 | D | -0.69 | -1.06 | 0.0 | -0.04 | -1.73 | -1.74 | -1.98 | -1.47 | -1.81 | -0.8 | -1.76 | -0.74 | -0.86 | -0.99 | -2.79 | -0.97 | -0.8 | -0.94 | -2.07 | -2.15 |
| 50 | I | 0.28 | 0.13 | 0.99 | 0.95 | -0.15 | -0.2 | -0.26 | 0.0 | -0.18 | 0.25 | -0.1 | 0.28 | 0.21 | 0.13 | -0.64 | 0.14 | 0.24 | 0.17 | -0.49 | -0.36 |
| 51 | E | -0.68 | -1.07 | 0.04 | 0.0 | -1.76 | -1.76 | -2.0 | -1.52 | -1.86 | -0.79 | -1.79 | -0.74 | -0.86 | -1.0 | -2.84 | -0.97 | -0.79 | -0.95 | -2.07 | -2.16 |
| 52 | Q | 0.16 | 0.0 | 0.95 | 0.89 | -0.28 | -0.31 | -0.4 | -0.1 | -0.33 | 0.12 | -0.24 | 0.16 | 0.08 | 0.0 | -0.97 | 0.02 | 0.12 | 0.05 | -0.6 | -0.52 |
| 53 | W | 0.86 | 0.57 | 1.85 | 1.79 | 0.28 | 0.26 | 0.15 | 0.44 | 0.21 | 0.74 | 0.29 | 0.78 | 0.69 | 0.61 | -0.32 | 0.63 | 0.73 | 0.64 | 0.0 | 0.08 |
| 54 | F | 0.48 | 0.31 | 1.38 | 1.31 | 0.0 | -0.03 | -0.12 | 0.21 | -0.06 | 0.44 | 0.06 | 0.48 | 0.4 | 0.32 | -0.53 | 0.34 | 0.44 | 0.36 | -0.32 | -0.2 |
| 55 | T | 0.05 | -0.1 | 0.87 | 0.83 | -0.39 | -0.43 | -0.54 | -0.2 | -0.49 | 0.01 | -0.37 | 0.05 | -0.02 | -0.1 | -1.16 | -0.08 | 0.0 | -0.05 | -0.71 | -0.64 |
| 56 | E | -0.7 | -1.11 | 0.04 | 0.0 | -1.72 | -1.73 | -2.07 | -1.46 | -1.96 | -0.82 | -1.74 | -0.76 | -0.91 | -1.04 | -2.92 | -1.01 | -0.82 | -0.99 | -2.08 | -2.15 |
| 57 | D | -0.74 | -1.17 | 0.0 | -0.04 | -1.8 | -1.82 | -2.16 | -1.52 | -2.06 | -0.88 | -1.81 | -0.81 | -0.97 | -1.1 | -3.01 | -1.08 | -0.88 | -1.05 | -2.17 | -2.22 |
| 58 | P | 0.08 | -0.08 | 0.94 | 0.9 | -0.35 | -0.38 | -0.52 | -0.17 | -0.48 | 0.04 | -0.32 | 0.07 | 0.0 | -0.07 | -1.15 | -0.06 | 0.04 | -0.03 | -0.68 | -0.61 |
| 59 | G | 0.58 | 0.38 | 1.62 | 1.55 | 0.1 | 0.0 | -0.07 | 0.29 | -0.04 | 0.53 | 0.14 | 0.57 | 0.48 | 0.4 | -0.51 | 0.41 | 0.52 | 0.44 | -0.23 | -0.12 |
| 60 | P | 0.08 | -0.08 | 0.97 | 0.93 | -0.34 | -0.38 | -0.54 | -0.17 | -0.52 | 0.04 | -0.32 | 0.07 | 0.0 | -0.07 | -1.2 | -0.06 | 0.04 | -0.03 | -0.68 | -0.6 |
| 61 | D | -0.82 | -1.29 | 0.0 | -0.04 | -1.92 | -1.94 | -2.35 | -1.62 | -2.3 | -1.0 | -1.93 | -0.91 | -1.09 | -1.22 | -3.24 | -1.19 | -1.0 | -1.16 | -2.3 | -2.35 |
| 62 | E | -0.8 | -1.29 | 0.04 | 0.0 | -1.9 | -1.91 | -2.36 | -1.61 | -2.33 | -1.0 | -1.91 | -0.91 | -1.08 | -1.22 | -3.26 | -1.18 | -1.0 | -1.16 | -2.27 | -2.33 |
| 63 | A | 0.0 | -0.16 | 0.87 | 0.83 | -0.48 | -0.51 | -0.72 | -0.29 | -0.74 | -0.04 | -0.47 | -0.01 | -0.08 | -0.14 | -1.6 | -0.13 | -0.05 | -0.11 | -0.77 | -0.71 |
| 64 | P | 0.08 | -0.08 | 1.05 | 0.99 | -0.34 | -0.38 | -0.6 | -0.17 | -1.0 | 0.04 | -0.32 | 0.07 | 0.0 | -0.07 | -1.8 | -0.06 | 0.04 | -0.03 | -0.68 | -0.6 |
| 65 | R | 1.36 | 1.13 | 2.82 | 2.73 | 0.78 | 0.72 | 0.51 | 1.03 | 0.5 | 1.31 | 0.84 | 1.36 | 1.26 | 1.15 | 0.0 | 1.17 | 1.31 | 1.21 | 0.33 | 0.49 |
| 66 | M | 0.38 | 0.24 | 1.5 | 1.42 | -0.04 | -0.1 | -0.25 | 0.15 | -0.52 | 0.36 | 0.0 | 0.39 | 0.32 | 0.24 | -1.21 | 0.26 | 0.36 | 0.29 | -0.39 | -0.26 |
| 67 | P | 0.07 | -0.08 | 1.07 | 1.01 | -0.34 | -0.38 | -0.62 | -0.17 | -0.67 | 0.04 | -0.32 | 0.07 | 0.0 | -0.07 | -1.47 | -0.06 | 0.04 | -0.03 | -0.68 | -0.6 |
| 68 | E | -0.71 | -1.23 | 0.04 | 0.0 | -1.92 | -1.94 | -2.79 | -1.56 | -2.89 | -0.95 | -1.92 | -0.87 | -1.03 | -1.16 | -3.74 | -1.13 | -0.96 | -1.1 | -2.37 | -2.45 |
| 69 | A | 0.0 | -0.16 | 0.92 | 0.87 | -0.45 | -0.48 | -0.92 | -0.26 | -1.07 | -0.04 | -0.44 | -0.01 | -0.08 | -0.14 | -1.92 | -0.13 | -0.04 | -0.11 | -0.77 | -0.69 |
| 70 | A | 0.0 | -0.16 | 0.92 | 0.87 | -0.44 | -0.48 | -1.1 | -0.25 | -1.29 | -0.04 | -0.43 | -0.01 | -0.08 | -0.14 | -2.09 | -0.13 | -0.04 | -0.11 | -0.77 | -0.68 |
| 71 | P | 0.07 | -0.08 | 0.94 | 0.89 | -0.32 | -0.37 | -1.1 | -0.16 | -1.26 | 0.04 | -0.28 | 0.07 | 0.0 | -0.07 | -1.94 | -0.05 | 0.04 | -0.03 | -0.68 | -0.55 |
| 72 | P | 0.07 | -0.08 | 0.91 | 0.86 | -0.32 | -0.37 | -1.28 | -0.16 | -1.46 | 0.04 | -0.28 | 0.07 | 0.0 | -0.07 | -2.07 | -0.05 | 0.03 | -0.03 | -0.68 | -0.55 |
| 73 | V | 0.11 | -0.04 | 0.98 | 0.93 | -0.28 | -0.34 | -1.32 | -0.11 | -1.48 | 0.08 | -0.24 | 0.11 | 0.04 | -0.03 | -2.08 | -0.02 | 0.08 | 0.0 | -0.63 | -0.5 |
| 74 | A | 0.0 | -0.15 | 0.83 | 0.79 | -0.43 | -0.46 | -1.65 | -0.23 | -1.86 | -0.04 | -0.41 | -0.01 | -0.07 | -0.14 | -2.37 | -0.13 | -0.04 | -0.1 | -0.77 | -0.67 |
| 75 | P | 0.07 | -0.08 | 1.02 | 0.98 | -0.32 | -0.36 | -1.38 | -0.15 | -1.55 | 0.04 | -0.28 | 0.07 | 0.0 | -0.07 | -2.1 | -0.05 | 0.03 | -0.03 | -0.68 | -0.55 |
| 76 | A | 0.0 | -0.15 | 0.96 | 0.93 | -0.42 | -0.46 | -1.52 | -0.22 | -1.74 | -0.04 | -0.4 | -0.01 | -0.07 | -0.14 | -2.24 | -0.13 | -0.04 | -0.1 | -0.76 | -0.67 |
| 77 | P | 0.06 | -0.08 | 1.2 | 1.16 | -0.32 | -0.36 | -1.2 | -0.15 | -1.4 | 0.04 | -0.27 | 0.07 | 0.0 | -0.07 | -1.86 | -0.05 | 0.03 | -0.03 | -0.68 | -0.54 |
| 78 | A | 0.0 | -0.15 | 1.13 | 1.1 | -0.41 | -0.45 | -1.31 | -0.22 | -1.52 | -0.04 | -0.39 | -0.01 | -0.07 | -0.14 | -1.97 | -0.12 | -0.04 | -0.1 | -0.76 | -0.66 |
| 79 | A | 0.0 | -0.15 | 1.22 | 1.2 | -0.41 | -0.45 | -1.18 | -0.22 | -1.41 | -0.04 | -0.39 | -0.01 | -0.07 | -0.13 | -1.82 | -0.12 | -0.04 | -0.1 | -0.75 | -0.65 |
| 80 | P | 0.06 | -0.08 | 1.48 | 1.46 | -0.31 | -0.36 | -0.85 | -0.15 | -1.06 | 0.04 | -0.27 | 0.06 | 0.0 | -0.07 | -1.43 | -0.05 | 0.03 | -0.03 | -0.67 | -0.54 |
| 81 | T | 0.04 | -0.1 | 1.57 | 1.55 | -0.33 | -0.38 | -0.74 | -0.17 | -0.96 | 0.02 | -0.28 | 0.04 | -0.02 | -0.08 | -1.3 | -0.07 | 0.0 | -0.05 | -0.7 | -0.56 |
| 82 | P | 0.06 | -0.08 | 1.51 | 1.49 | -0.31 | -0.36 | -0.86 | -0.15 | -1.09 | 0.04 | -0.26 | 0.06 | 0.0 | -0.07 | -1.46 | -0.05 | 0.03 | -0.03 | -0.67 | -0.54 |
| 83 | A | 0.0 | -0.14 | 1.28 | 1.26 | -0.4 | -0.44 | -1.21 | -0.21 | -1.47 | -0.03 | -0.38 | -0.0 | -0.06 | -0.13 | -1.88 | -0.11 | -0.04 | -0.1 | -0.74 | -0.64 |
| 84 | A | 0.0 | -0.14 | 1.23 | 1.2 | -0.39 | -0.43 | -1.35 | -0.21 | -1.61 | -0.03 | -0.37 | -0.0 | -0.06 | -0.13 | -2.07 | -0.11 | -0.04 | -0.09 | -0.74 | -0.63 |
| 85 | P | 0.05 | -0.08 | 1.31 | 1.27 | -0.31 | -0.35 | -1.27 | -0.14 | -1.55 | 0.03 | -0.26 | 0.05 | 0.0 | -0.07 | -2.02 | -0.05 | 0.02 | -0.03 | -0.66 | -0.53 |
| 86 | A | 0.0 | -0.14 | 1.12 | 1.09 | -0.38 | -0.43 | -1.58 | -0.2 | -1.94 | -0.02 | -0.36 | -0.0 | -0.05 | -0.12 | -2.39 | -0.11 | -0.03 | -0.09 | -0.72 | -0.62 |
| 87 | P | 0.04 | -0.08 | 1.19 | 1.15 | -0.3 | -0.35 | -1.48 | -0.14 | -1.83 | 0.03 | -0.25 | 0.05 | 0.0 | -0.07 | -2.33 | -0.05 | 0.02 | -0.03 | -0.66 | -0.52 |
| 88 | A | 0.0 | -0.13 | 1.06 | 1.02 | -0.38 | -0.42 | -1.76 | -0.2 | -2.14 | -0.02 | -0.35 | 0.0 | -0.04 | -0.11 | -2.59 | -0.1 | -0.03 | -0.08 | -0.7 | -0.62 |
| 89 | P | 0.04 | -0.08 | 1.14 | 1.09 | -0.3 | -0.35 | -1.54 | -0.14 | -1.96 | 0.02 | -0.25 | 0.04 | 0.0 | -0.07 | -2.47 | -0.05 | 0.02 | -0.04 | -0.64 | -0.52 |
| 90 | S | 0.09 | -0.02 | 1.25 | 1.2 | -0.25 | -0.3 | -1.39 | -0.09 | -1.85 | 0.08 | -0.2 | 0.1 | 0.06 | 0.0 | -2.44 | 0.0 | 0.08 | 0.02 | -0.58 | -0.47 |
| 91 | W | 0.72 | 0.49 | 2.18 | 2.13 | 0.3 | 0.27 | -0.91 | 0.43 | -1.33 | 0.64 | 0.33 | 0.65 | 0.62 | 0.53 | -1.77 | 0.56 | 0.62 | 0.56 | 0.0 | 0.11 |
| 92 | P | 0.01 | -0.08 | 1.3 | 1.24 | -0.28 | -0.34 | -1.1 | -0.14 | -1.71 | 0.02 | -0.23 | 0.02 | 0.0 | -0.07 | -2.44 | -0.05 | 0.01 | -0.04 | -0.64 | -0.52 |
| 93 | L | -0.01 | -0.1 | 1.34 | 1.27 | -0.29 | -0.34 | -1.0 | -0.14 | -1.64 | 0.0 | -0.23 | 0.01 | -0.01 | -0.08 | -2.36 | -0.07 | -0.01 | -0.05 | -0.65 | -0.51 |
