## Supplementary material for "Accelerated Missense Mutation Identification in Intrinsically Disordered Proteins using Deep Learning": Mutation heat-map ProTaC

| ID | WT | A | C | D | E | F | G | H | I | K | L | M | N | P | Q | R | S | T | V | W | Y |
|---|---|---|---|---|---|---|---|---|---|---|---|---|---|---|---|---|---|---|---|---|---|
| 1 | M | 0.0 | 0.0 | 0.0 | 0.0 | 0.0 | 0.0 | 0.0 | -0.08 | 0.0 | 0.0 | 0.0 | 0.0 | 0.0 | 0.0 | 0.0 | 0.0 | 0.0 | 0.0 | 0.0 | 0.0 |
| 2 | A | 0.0 | -3.04 | 0.0 | 0.0 | -2.84 | -0.39 | -5.41 | -4.08 | -6.79 | -2.61 | -4.1 | -1.99 | -2.9 | -2.53 | -6.76 | -2.26 | -2.2 | -2.52 | 0.0 | -1.86 |
| 3 | H | 0.0 | 0.0 | 0.0 | 0.0 | 0.0 | 0.0 | 0.0 | 0.0 | -3.66 | 0.0 | 0.0 | 0.0 | 0.0 | 0.0 | -3.39 | 0.0 | 0.0 | 0.0 | 0.0 | 0.0 |
| 4 | H | 0.0 | 0.0 | 0.0 | 0.0 | 0.0 | 0.0 | 0.0 | 0.0 | -2.99 | 0.0 | 0.0 | 0.0 | 0.0 | 0.0 | -2.76 | 0.0 | 0.0 | 0.0 | 0.0 | 0.0 |
| 5 | H | 0.0 | 0.0 | 0.31 | 0.35 | 0.0 | 0.0 | 0.0 | 0.0 | -2.18 | 0.0 | 0.0 | 0.0 | 0.0 | 0.0 | -2.0 | 0.0 | 0.0 | 0.0 | 0.0 | 0.0 |
| 6 | H | 0.0 | 0.0 | 0.31 | 0.34 | 0.0 | 0.0 | 0.0 | 0.0 | -2.09 | 0.0 | 0.0 | 0.0 | 0.0 | 0.0 | -1.97 | 0.0 | 0.0 | 0.0 | 0.0 | 0.0 |
| 7 | H | 0.0 | 0.0 | 0.0 | 0.0 | 0.0 | 0.0 | 0.0 | 0.0 | -2.71 | 0.0 | 0.0 | 0.0 | 0.0 | 0.0 | -2.67 | 0.0 | 0.0 | 0.0 | 0.0 | 0.0 |
| 8 | H | 0.0 | 0.0 | 0.0 | 0.0 | 0.0 | 0.0 | 0.0 | 0.0 | -3.18 | 0.0 | 0.0 | 0.0 | 0.0 | 0.0 | -3.25 | 0.0 | 0.0 | 0.0 | 0.0 | 0.0 |
| 9 | S | 0.0 | -0.28 | 0.0 | 0.0 | -0.19 | 0.0 | -4.35 | -1.28 | -5.69 | 0.0 | -1.54 | 0.0 | -0.19 | 0.0 | -5.7 | 0.0 | 0.0 | 0.0 | 0.0 | 0.0 |
| 10 | A | 0.0 | -3.36 | 0.0 | 0.0 | -3.29 | -0.79 | -5.38 | -4.23 | -6.62 | -2.88 | -4.34 | -2.12 | -3.27 | -2.91 | -6.65 | -2.64 | -2.42 | -2.86 | 0.0 | -2.39 |
| 11 | A | 0.0 | -3.39 | 0.0 | 0.0 | -3.34 | -0.84 | -5.37 | -4.25 | -6.59 | -2.9 | -4.37 | -2.13 | -3.3 | -2.93 | -6.63 | -2.67 | -2.43 | -2.88 | 0.0 | -2.45 |
| 12 | L | 0.0 | -0.04 | 0.0 | 0.0 | -0.03 | 0.0 | -3.82 | -1.08 | -5.43 | 0.0 | -1.38 | 0.0 | 0.0 | 0.0 | -5.5 | 0.0 | 0.0 | 0.0 | 0.0 | 0.0 |
| 13 | E | 0.0 | 0.0 | -0.2 | 0.0 | 0.0 | 0.0 | -7.65 | 0.0 | -8.02 | 0.0 | 0.0 | 0.0 | 0.0 | 0.0 | -8.03 | 0.0 | 0.0 | 0.0 | 0.0 | 0.0 |
| 14 | V | 0.0 | -0.07 | 0.0 | 0.0 | -0.17 | 0.0 | -3.81 | -1.22 | -5.38 | 0.0 | -1.58 | 0.0 | -0.04 | 0.0 | -5.49 | 0.0 | 0.0 | 0.0 | 0.0 | 0.0 |
| 15 | L | 0.0 | -0.04 | 0.0 | 0.0 | -0.28 | 0.0 | -3.71 | -1.33 | -5.34 | 0.0 | -1.68 | 0.0 | -0.0 | 0.0 | -5.5 | 0.0 | 0.0 | 0.0 | 0.0 | 0.0 |
| 16 | F | 0.0 | 0.0 | 0.0 | 0.0 | 0.0 | 0.0 | -2.81 | -0.19 | -5.04 | 0.0 | -0.53 | 0.0 | 0.0 | 0.0 | -5.15 | 0.0 | 0.0 | 0.0 | 0.0 | 0.0 |
| 17 | Q | 0.0 | -0.01 | 0.0 | 0.0 | -0.38 | 0.0 | -3.59 | -1.44 | -5.26 | 0.0 | -1.82 | 0.0 | 0.0 | 0.0 | -5.48 | 0.0 | 0.0 | 0.0 | 0.0 | 0.0 |
| 18 | G | 0.0 | -1.99 | 0.0 | 0.0 | -2.24 | 0.0 | -4.77 | -3.22 | -5.95 | -1.47 | -3.6 | -0.61 | -1.92 | -1.55 | -6.13 | -1.26 | -0.95 | -1.46 | 0.0 | -1.38 |
| 19 | P | 0.0 | 0.0 | 0.0 | 0.0 | -0.01 | 0.0 | -3.07 | -0.98 | -5.04 | 0.0 | -1.39 | 0.0 | 0.0 | 0.0 | -5.28 | 0.0 | 0.0 | 0.0 | 0.0 | 0.0 |
| 20 | M | 0.0 | 0.0 | 0.0 | 0.0 | 0.0 | 0.0 | -1.47 | 0.0 | -4.36 | 0.0 | 0.0 | 0.0 | 0.0 | 0.0 | -4.54 | 0.0 | 0.0 | 0.0 | 0.0 | 0.0 |
| 21 | S | 0.0 | -0.25 | 0.0 | 0.0 | -0.6 | 0.0 | -3.71 | -1.62 | -5.21 | 0.0 | -2.03 | 0.0 | -0.18 | 0.0 | -5.41 | 0.0 | 0.0 | 0.0 | 0.0 | 0.0 |
| 22 | D | 0.0 | 0.0 | 0.0 | 0.0 | 0.0 | 0.0 | -7.35 | 0.0 | -7.82 | 0.0 | 0.0 | 0.0 | 0.0 | 0.0 | -7.87 | 0.0 | 0.0 | 0.0 | 0.0 | 0.0 |
| 23 | A | 0.0 | -3.55 | 0.0 | 0.0 | -3.74 | -1.17 | -5.24 | -4.39 | -6.26 | -3.04 | -4.56 | -2.2 | -3.48 | -3.13 | -6.4 | -2.85 | -2.54 | -3.05 | 0.0 | -2.88 |
| 24 | A | 0.0 | -3.56 | 0.0 | 0.0 | -3.74 | -1.16 | -5.23 | -4.38 | -6.24 | -3.05 | -4.56 | -2.21 | -3.49 | -3.14 | -6.39 | -2.86 | -2.55 | -3.06 | 0.0 | -2.89 |
| 25 | V | 0.0 | -0.06 | 0.0 | 0.0 | -0.25 | 0.0 | -3.33 | -1.24 | -5.01 | 0.0 | -1.69 | 0.0 | -0.04 | 0.0 | -5.16 | 0.0 | 0.0 | 0.0 | 0.0 | 0.0 |
| 26 | D | -5.17 | -6.47 | 0.0 | 0.0 | -6.48 | -5.62 | -7.24 | -6.78 | -7.72 | -6.33 | -6.88 | -6.04 | -6.45 | -6.34 | -7.78 | -6.26 | -6.16 | -6.33 | -4.62 | -6.18 |
| 27 | T | 0.0 | -0.58 | 0.0 | 0.0 | -0.79 | 0.0 | -3.75 | -1.79 | -5.12 | -0.1 | -2.24 | 0.0 | -0.56 | -0.19 | -5.28 | 0.0 | 0.0 | -0.11 | 0.0 | -0.02 |
| 28 | S | 0.0 | -0.24 | 0.0 | 0.0 | -0.49 | 0.0 | -3.43 | -1.47 | -5.0 | 0.0 | -1.92 | 0.0 | -0.18 | 0.0 | -5.17 | 0.0 | 0.0 | 0.0 | 0.0 | 0.0 |
| 29 | S | 0.0 | -0.24 | 0.0 | 0.0 | -0.54 | 0.0 | -3.37 | -1.52 | -4.95 | 0.0 | -1.98 | 0.0 | -0.18 | 0.0 | -5.15 | 0.0 | 0.0 | 0.0 | 0.0 | 0.0 |
| 30 | E | -5.25 | -6.55 | -0.18 | 0.0 | -6.64 | -5.82 | -7.27 | -6.95 | -7.72 | -6.41 | -7.03 | -6.14 | -6.53 | -6.43 | -7.82 | -6.36 | -6.26 | -6.42 | -4.77 | -6.38 |
| 31 | I | 0.0 | 0.0 | 0.0 | 0.0 | 0.0 | 0.0 | -1.1 | 0.0 | -3.69 | 0.0 | 0.0 | 0.0 | 0.0 | 0.0 | -4.15 | 0.0 | 0.0 | 0.0 | 0.0 | 0.0 |
| 32 | T | 0.0 | -0.59 | 0.0 | 0.0 | -1.06 | 0.0 | -3.6 | -2.04 | -5.0 | -0.1 | -2.51 | 0.0 | -0.58 | -0.21 | -5.25 | 0.0 | 0.0 | -0.12 | 0.0 | -0.26 |
| 33 | T | 0.0 | -0.6 | 0.0 | 0.0 | -0.92 | 0.0 | -3.59 | -1.9 | -4.98 | -0.1 | -2.38 | 0.0 | -0.58 | -0.21 | -5.2 | 0.0 | 0.0 | -0.13 | 0.0 | -0.13 |
| 34 | K | 0.0 | 0.0 | 0.0 | 0.0 | 0.0 | 0.0 | 0.0 | 0.0 | 0.0 | 0.0 | 0.0 | 0.0 | 0.0 | 0.0 | -0.34 | 0.0 | 0.0 | 0.0 | 0.0 | 0.0 |
| 35 | D | -4.93 | -6.3 | 0.0 | 0.0 | -6.34 | -5.44 | -7.03 | -6.63 | -7.54 | -6.12 | -6.74 | -5.82 | -6.28 | -6.14 | -7.62 | -6.05 | -5.94 | -6.12 | -4.4 | -6.01 |
| 36 | L | 0.0 | -0.05 | 0.0 | 0.0 | -0.31 | 0.0 | -2.9 | -1.26 | -4.61 | 0.0 | -1.75 | 0.0 | -0.03 | 0.0 | -4.86 | 0.0 | 0.0 | 0.0 | 0.0 | 0.0 |
| 37 | K | 0.0 | 0.0 | 0.0 | 0.0 | 0.0 | 0.0 | 0.0 | 0.0 | 0.0 | 0.0 | 0.0 | 0.0 | 0.0 | 0.0 | -0.32 | 0.0 | 0.0 | 0.0 | 0.0 | 0.0 |
| 38 | E | -5.07 | -6.41 | -0.18 | 0.0 | -6.44 | -5.56 | -7.09 | -6.73 | -8.66 | -6.26 | -6.85 | -5.95 | -6.4 | -6.28 | -8.69 | -6.19 | -6.08 | -6.26 | -4.53 | -6.13 |
| 39 | K | 0.0 | 0.0 | 0.0 | 0.0 | 0.0 | 0.0 | 0.0 | 0.0 | 0.0 | 0.0 | 0.0 | 0.0 | 0.0 | 0.0 | -0.33 | 0.0 | 0.0 | 0.0 | 0.0 | 0.0 |
| 40 | K | 0.0 | 0.0 | 0.0 | 0.0 | 0.0 | 0.0 | 0.0 | 0.0 | 0.0 | 0.0 | 0.0 | 0.0 | 0.0 | 0.0 | -0.34 | 0.0 | 0.0 | 0.0 | 0.0 | 0.0 |
| 41 | E | -4.94 | -6.32 | -0.18 | 0.0 | -6.35 | -5.46 | -6.99 | -6.64 | -8.21 | -6.14 | -6.76 | -5.83 | -6.3 | -6.16 | -8.29 | -6.07 | -5.96 | -6.14 | -4.4 | -6.03 |
| 42 | V | 0.0 | -0.06 | 0.0 | 0.0 | -0.35 | 0.0 | -2.65 | -1.28 | -4.36 | 0.0 | -1.82 | 0.0 | -0.05 | 0.0 | -4.63 | 0.0 | 0.0 | 0.0 | 0.0 | 0.0 |
| 43 | V | 0.0 | -0.06 | 0.0 | 0.0 | -0.36 | 0.0 | -2.6 | -1.28 | -4.3 | 0.0 | -1.83 | 0.0 | -0.05 | 0.0 | -4.58 | 0.0 | 0.0 | 0.0 | 0.0 | 0.0 |
| 44 | E | -4.76 | -6.18 | -0.17 | 0.0 | -6.24 | -5.34 | -6.85 | -6.54 | -7.27 | -6.0 | -6.66 | -5.69 | -6.16 | -6.03 | -7.43 | -5.94 | -5.82 | -6.0 | -4.25 | -5.9 |
| 45 | E | -4.72 | -6.14 | -0.17 | 0.0 | -6.21 | -5.31 | -6.81 | -6.52 | -7.23 | -5.96 | -6.65 | -5.66 | -6.13 | -5.99 | -7.39 | -5.9 | -5.78 | -5.97 | -4.22 | -5.88 |
| 46 | A | 0.0 | -3.7 | 0.0 | 0.0 | -4.07 | -1.4 | -4.95 | -4.5 | -5.56 | -3.18 | -4.72 | -2.28 | -3.65 | -3.31 | -5.8 | -3.02 | -2.65 | -3.2 | 0.0 | -3.25 |
| 47 | E | -4.66 | -6.1 | -0.17 | 0.0 | -6.2 | -5.3 | -6.76 | -6.52 | -7.17 | -5.92 | -6.64 | -5.61 | -6.08 | -5.95 | -7.35 | -5.86 | -5.74 | -5.93 | -4.17 | -5.87 |
| 48 | N | 0.0 | -1.01 | 0.0 | 0.0 | -1.58 | 0.0 | -3.29 | -2.49 | -4.5 | -0.5 | -3.03 | 0.0 | -1.0 | -0.64 | -4.89 | -0.37 | -0.01 | -0.54 | 0.0 | -0.79 |
| 49 | G | 0.0 | -1.79 | 0.0 | 0.0 | -2.36 | 0.0 | -4.11 | -3.23 | -4.89 | -1.25 | -3.77 | -0.32 | -1.74 | -1.4 | -5.23 | -1.08 | -0.7 | -1.27 | 0.0 | -1.58 |
| 50 | R | 0.0 | 0.0 | 0.0 | 0.0 | 0.0 | 0.0 | 0.0 | 0.0 | 0.0 | 0.0 | 0.0 | 0.0 | 0.0 | 0.0 | 0.0 | 0.0 | 0.0 | 0.0 | 0.0 | 0.0 |
| 51 | D | -1.03 | -4.49 | 0.0 | 0.0 | -4.7 | -2.83 | -6.5 | -5.1 | -6.94 | -4.27 | -5.26 | -3.6 | -4.47 | -4.32 | -7.14 | -4.2 | -3.97 | -4.28 | -0.14 | -4.32 |
| 52 | A | 0.0 | -3.72 | 0.0 | 0.0 | -4.15 | -1.51 | -4.87 | -4.55 | -5.42 | -3.2 | -4.79 | -2.3 | -3.68 | -3.35 | -5.7 | -3.04 | -2.67 | -3.23 | 0.0 | -3.4 |
| 53 | P | 0.0 | 0.0 | 0.0 | 0.0 | 0.0 | 0.0 | -1.65 | -0.83 | -3.16 | 0.0 | -1.39 | 0.0 | 0.0 | 0.0 | -3.96 | 0.0 | 0.0 | 0.0 | 0.0 | 0.0 |
| 54 | A | 0.0 | -3.73 | 0.0 | 0.0 | -4.16 | -1.53 | -4.84 | -4.56 | -5.37 | -3.21 | -4.8 | -2.3 | -3.69 | -3.36 | -5.66 | -3.05 | -2.68 | -3.23 | 0.0 | -3.42 |
| 55 | N | 0.0 | -1.02 | 0.0 | 0.0 | -1.63 | 0.0 | -3.03 | -2.53 | -4.26 | -0.51 | -3.09 | 0.0 | -1.01 | -0.66 | -4.67 | -0.38 | -0.01 | -0.55 | 0.0 | -0.85 |
| 56 | G | 0.0 | -1.98 | 0.0 | 0.0 | -2.38 | 0.0 | -4.0 | -3.24 | -4.67 | -1.44 | -3.8 | -0.47 | -1.94 | -1.6 | -5.02 | -1.27 | -0.86 | -1.46 | 0.0 | -1.61 |
| 57 | N | 0.0 | -1.02 | 0.0 | 0.0 | -1.64 | 0.0 | -2.94 | -2.53 | -4.18 | -0.52 | -3.1 | 0.0 | -1.02 | -0.66 | -4.59 | -0.38 | -0.01 | -0.55 | 0.0 | -0.86 |
| 58 | A | 0.0 | -3.74 | 0.0 | 0.0 | -4.17 | -1.54 | -4.76 | -4.56 | -5.24 | -3.22 | -4.81 | -2.31 | -3.7 | -3.38 | -5.53 | -3.07 | -2.68 | -3.25 | 0.0 | -3.44 |
| 59 | N | 0.0 | -1.02 | 0.0 | 0.0 | -1.47 | 0.0 | -2.82 | -2.36 | -4.0 | -0.52 | -2.92 | 0.0 | -1.02 | -0.66 | -4.41 | -0.38 | -0.01 | -0.55 | 0.0 | -0.7 |
| 60 | E | -0.64 | -4.34 | -0.16 | 0.0 | -4.48 | -2.37 | -6.26 | -4.91 | -6.59 | -4.11 | -5.1 | -3.23 | -4.32 | -4.17 | -6.78 | -3.97 | -3.6 | -4.13 | 0.0 | -4.1 |
| 61 | E | -4.03 | -5.65 | -0.16 | 0.0 | -5.78 | -4.85 | -6.21 | -6.13 | -6.54 | -5.48 | -6.31 | -5.17 | -5.64 | -5.52 | -6.74 | -5.42 | -5.3 | -5.49 | -3.1 | -5.48 |
| 62 | N | 0.0 | -1.02 | 0.0 | 0.0 | -1.67 | 0.0 | -2.68 | -2.54 | -3.7 | -0.52 | -3.12 | 0.0 | -1.02 | -0.67 | -4.36 | -0.39 | -0.01 | -0.56 | 0.0 | -0.89 |
| 63 | G | 0.0 | -1.99 | 0.0 | 0.0 | -2.41 | 0.0 | -3.64 | -3.25 | -4.34 | -1.45 | -3.82 | -0.48 | -1.95 | -1.63 | -4.7 | -1.28 | -0.87 | -1.47 | 0.0 | -1.64 |
| 64 | E | -3.79 | -5.57 | -0.16 | 0.0 | -5.76 | -4.81 | -6.08 | -6.11 | -6.41 | -5.4 | -6.29 | -5.09 | -5.56 | -5.44 | -6.66 | -5.34 | -5.22 | -5.41 | -2.99 | -5.46 |
| 65 | Q | 0.0 | 0.0 | 0.0 | 0.0 | -0.38 | 0.0 | -1.43 | -1.27 | -2.31 | 0.0 | -1.85 | 0.0 | 0.0 | 0.0 | -3.16 | 0.0 | 0.0 | 0.0 | 0.0 | 0.0 |
| 66 | E | -3.61 | -5.51 | -0.16 | 0.0 | -5.62 | -4.64 | -5.99 | -5.96 | -6.3 | -5.34 | -6.14 | -5.04 | -5.5 | -5.38 | -6.5 | -5.28 | -5.16 | -5.35 | -2.74 | -5.33 |
| 67 | A | 0.0 | -3.77 | 0.0 | 0.0 | -4.16 | -1.48 | -4.58 | -4.54 | -4.92 | -3.25 | -4.79 | -2.33 | -3.73 | -3.41 | -5.22 | -3.09 | -2.7 | -3.27 | 0.0 | -3.42 |
| 68 | D | -2.9 | -5.27 | 0.0 | 0.0 | -5.38 | -4.31 | -5.72 | -5.7 | -5.99 | -5.1 | -5.88 | -4.73 | -5.26 | -5.14 | -6.23 | -5.04 | -4.89 | -5.11 | -2.17 | -5.1 |
| 69 | N | 0.0 | -1.03 | 0.0 | 0.0 | -1.44 | 0.0 | -2.39 | -2.32 | -3.07 | -0.52 | -2.89 | 0.0 | -1.03 | -0.68 | -3.86 | -0.39 | -0.01 | -0.56 | 0.0 | -0.68 |
| 70 | E | -3.28 | -5.41 | -0.16 | 0.0 | -5.5 | -4.49 | -5.82 | -5.83 | -6.08 | -5.23 | -6.02 | -4.9 | -5.39 | -5.28 | -6.32 | -5.18 | -5.06 | -5.24 | -2.46 | -5.21 |
| 71 | V | 0.0 | -0.05 | 0.0 | 0.0 | -0.48 | 0.0 | -1.38 | -1.35 | -2.01 | 0.0 | -1.96 | 0.0 | -0.05 | 0.0 | -2.81 | 0.0 | 0.0 | 0.0 | 0.0 | 0.0 |
| 72 | D | -2.56 | -5.16 | 0.0 | 0.0 | -5.26 | -4.17 | -5.55 | -5.58 | -5.75 | -4.98 | -5.76 | -4.58 | -5.15 | -5.03 | -6.0 | -4.91 | -4.74 | -4.99 | -1.87 | -4.99 |
| 73 | E | -3.0 | -5.31 | -0.16 | 0.0 | -5.42 | -4.37 | -5.68 | -5.74 | -5.88 | -5.14 | -5.92 | -4.78 | -5.3 | -5.18 | -6.13 | -5.08 | -4.94 | -5.15 | -2.21 | -5.13 |
| 74 | E | -2.97 | -5.3 | -0.16 | 0.0 | -5.42 | -4.36 | -5.67 | -5.74 | -5.86 | -5.13 | -5.92 | -4.76 | -5.29 | -5.17 | -6.12 | -5.08 | -4.93 | -5.14 | -2.19 | -5.13 |
| 75 | C | 0.0 | 0.0 | 0.0 | 0.0 | 0.0 | 0.0 | -0.34 | -0.47 | -0.84 | 0.0 | -1.06 | 0.0 | 0.0 | 0.0 | -1.72 | 0.0 | 0.0 | 0.0 | 0.0 | 0.0 |
| 76 | E | -2.87 | -5.27 | -0.16 | 0.0 | -5.41 | -4.36 | -5.62 | -5.73 | -5.79 | -5.1 | -5.92 | -4.72 | -5.26 | -5.14 | -6.08 | -5.04 | -4.89 | -5.11 | -2.1 | -5.13 |
| 77 | E | -2.83 | -5.26 | -0.16 | 0.0 | -5.46 | -4.4 | -5.6 | -5.78 | -5.76 | -5.08 | -5.97 | -4.7 | -5.24 | -5.13 | -6.11 | -5.03 | -4.87 | -5.09 | -2.15 | -5.17 |
| 78 | G | 0.0 | -1.78 | 0.0 | 0.0 | -2.45 | 0.0 | -2.92 | -3.27 | -3.39 | -1.24 | -3.87 | -0.29 | -1.74 | -1.43 | -4.26 | -1.07 | -0.67 | -1.26 | 0.0 | -1.7 |
| 79 | G | 0.0 | -1.78 | -5.53 | -5.35 | -2.45 | 0.0 | -2.9 | -3.27 | -3.34 | -1.24 | -3.87 | -0.29 | -1.73 | -1.43 | -4.24 | -1.07 | -0.67 | -1.26 | 0.0 | -1.71 |
| 80 | E | 0.0 | 0.0 | -0.15 | 0.0 | 0.0 | 0.0 | 0.0 | 0.0 | 0.0 | 0.0 | 0.0 | 0.0 | 0.0 | 0.0 | 0.0 | 0.0 | 0.0 | 0.0 | 0.0 | 0.0 |
| 81 | E | 0.0 | 0.0 | -0.15 | 0.0 | 0.0 | 0.0 | 0.0 | 0.0 | 0.0 | 0.0 | 0.0 | 0.0 | 0.0 | 0.0 | 0.0 | 0.0 | 0.0 | 0.0 | 0.0 | 0.0 |
| 82 | E | 0.0 | 0.0 | -0.15 | 0.0 | 0.0 | 0.0 | 0.0 | 0.0 | 0.0 | 0.0 | 0.0 | 0.0 | 0.0 | 0.0 | 0.0 | 0.0 | 0.0 | 0.0 | 0.0 | 0.0 |
| 83 | E | 0.0 | 0.0 | -0.15 | 0.0 | 0.0 | 0.0 | 0.0 | 0.0 | 0.0 | 0.0 | 0.0 | 0.0 | 0.0 | 0.0 | 0.0 | 0.0 | 0.0 | 0.0 | 0.0 | 0.0 |
| 84 | E | 0.0 | 0.0 | -0.15 | 0.0 | 0.0 | 0.0 | 0.0 | 0.0 | 0.0 | 0.0 | 0.0 | 0.0 | 0.0 | 0.0 | 0.0 | 0.0 | 0.0 | 0.0 | 0.0 | 0.0 |
| 85 | E | 0.0 | 0.0 | -0.15 | 0.0 | 0.0 | 0.0 | 0.0 | 0.0 | 0.0 | 0.0 | 0.0 | 0.0 | 0.0 | 0.0 | 0.0 | 0.0 | 0.0 | 0.0 | 0.0 | 0.0 |
| 86 | E | 0.0 | 0.0 | -0.15 | 0.0 | 0.0 | 0.0 | 0.0 | 0.0 | 0.0 | 0.0 | 0.0 | 0.0 | 0.0 | 0.0 | 0.0 | 0.0 | 0.0 | 0.0 | 0.0 | 0.0 |
| 87 | E | 0.0 | 0.0 | -0.15 | 0.0 | 0.0 | 0.0 | 0.0 | 0.0 | 0.0 | 0.0 | 0.0 | 0.0 | 0.0 | 0.0 | 0.0 | 0.0 | 0.0 | 0.0 | 0.0 | 0.0 |
| 88 | G | 0.0 | -1.93 | -8.22 | -8.18 | -2.48 | 0.0 | -2.96 | -3.29 | -3.3 | -1.38 | -3.89 | -0.42 | -1.88 | -1.58 | -4.18 | -1.2 | -0.8 | -1.4 | 0.0 | -1.74 |
| 89 | D | -2.12 | -5.01 | 0.0 | 0.0 | -5.28 | -4.14 | -5.34 | -5.59 | -5.46 | -4.79 | -5.78 | -4.39 | -4.99 | -4.85 | -5.86 | -4.71 | -4.55 | -4.8 | -1.67 | -5.01 |
| 90 | G | 0.0 | -1.92 | 0.0 | 0.0 | -2.48 | 0.0 | -2.97 | -3.29 | -3.34 | -1.38 | -3.89 | -0.42 | -1.87 | -1.58 | -4.2 | -1.2 | -0.8 | -1.4 | 0.0 | -1.74 |
| 91 | E | -2.64 | -5.19 | -0.15 | 0.0 | -5.41 | -4.32 | -5.52 | -5.72 | -5.65 | -5.02 | -5.92 | -4.62 | -5.18 | -5.07 | -6.0 | -4.95 | -4.79 | -5.03 | -2.03 | -5.12 |
| 92 | E | -2.62 | -5.19 | -0.15 | 0.0 | -5.35 | -4.26 | -5.51 | -5.67 | -5.64 | -5.02 | -5.86 | -4.62 | -5.17 | -5.07 | -5.94 | -4.94 | -4.78 | -5.03 | -1.92 | -5.07 |
| 93 | E | -2.65 | -5.2 | -0.15 | 0.0 | -5.36 | -4.27 | -5.52 | -5.68 | -5.66 | -5.03 | -5.87 | -4.63 | -5.18 | -5.08 | -5.96 | -4.96 | -4.8 | -5.04 | -1.95 | -5.08 |
| 94 | D | -2.2 | -5.04 | 0.0 | 0.0 | -5.26 | -4.11 | -5.38 | -5.56 | -5.52 | -4.82 | -5.75 | -4.43 | -5.03 | -4.89 | -5.87 | -4.75 | -4.59 | -4.84 | -1.63 | -4.98 |
| 95 | G | 0.0 | -2.02 | -8.22 | -8.18 | -2.5 | 0.0 | -3.15 | -3.3 | -3.59 | -1.48 | -3.9 | -0.52 | -1.97 | -1.68 | -4.27 | -1.29 | -0.89 | -1.49 | 0.0 | -1.76 |
| 96 | D | -2.28 | -5.07 | 0.0 | 0.0 | -5.28 | -4.14 | -5.42 | -5.59 | -5.59 | -4.86 | -5.78 | -4.47 | -5.06 | -4.93 | -5.92 | -4.78 | -4.63 | -4.87 | -1.71 | -5.01 |
| 97 | E | -2.82 | -5.26 | -0.15 | 0.0 | -5.41 | -4.32 | -5.61 | -5.72 | -5.78 | -5.09 | -5.91 | -4.71 | -5.24 | -5.13 | -6.07 | -5.03 | -4.87 | -5.09 | -2.11 | -5.12 |
| 98 | D | -2.36 | -5.09 | 0.0 | 0.0 | -5.23 | -4.06 | -5.46 | -5.54 | -5.64 | -4.9 | -5.72 | -4.5 | -5.08 | -4.96 | -5.91 | -4.82 | -4.66 | -4.9 | -1.74 | -4.94 |
| 99 | E | -2.96 | -5.3 | -0.15 | 0.0 | -5.42 | -4.35 | -5.68 | -5.74 | -5.88 | -5.13 | -5.93 | -4.77 | -5.28 | -5.18 | -6.14 | -5.07 | -4.93 | -5.14 | -2.24 | -5.14 |
| 100 | E | -3.07 | -5.34 | -0.15 | 0.0 | -5.46 | -4.39 | -5.74 | -5.78 | -5.96 | -5.17 | -5.96 | -4.82 | -5.32 | -5.22 | -6.22 | -5.11 | -4.98 | -5.17 | -2.35 | -5.17 |
| 101 | A | 0.0 | -3.8 | 0.0 | 0.0 | -4.18 | -1.44 | -4.54 | -4.55 | -4.82 | -3.29 | -4.81 | -2.37 | -3.75 | -3.46 | -5.15 | -3.09 | -2.73 | -3.29 | 0.0 | -3.5 |
| 102 | E | -3.29 | -5.41 | -0.15 | 0.0 | -5.53 | -4.48 | -5.86 | -5.86 | -6.12 | -5.24 | -6.04 | -4.92 | -5.4 | -5.29 | -6.36 | -5.18 | -5.06 | -5.25 | -2.54 | -5.25 |
| 103 | S | 0.0 | -0.25 | 0.0 | 0.0 | -0.75 | 0.0 | -1.77 | -1.59 | -2.53 | 0.0 | -2.18 | 0.0 | -0.2 | 0.0 | -3.34 | 0.0 | 0.0 | 0.0 | 0.0 | -0.04 |
| 104 | A | 0.0 | -3.79 | 0.0 | 0.0 | -4.22 | -1.5 | -4.62 | -4.58 | -4.99 | -3.28 | -4.84 | -2.37 | -3.74 | -3.45 | -5.3 | -3.08 | -2.72 | -3.28 | 0.0 | -3.57 |
| 105 | T | 0.0 | -0.63 | 0.0 | 0.0 | -1.31 | 0.0 | -2.22 | -2.15 | -3.1 | -0.14 | -2.75 | 0.0 | -0.61 | -0.31 | -4.09 | -0.0 | 0.0 | -0.15 | 0.0 | -0.57 |
| 106 | G | 0.0 | -1.99 | 0.0 | 0.0 | -2.52 | 0.0 | -3.64 | -3.33 | -6.61 | -1.47 | -3.91 | -0.53 | -1.94 | -1.65 | -6.86 | -1.26 | -0.89 | -1.46 | 0.0 | -1.78 |
| 107 | K | 0.0 | 0.0 | 0.0 | 0.0 | 0.0 | 0.0 | 0.0 | 0.0 | 0.0 | 0.0 | 0.0 | 0.0 | 0.0 | 0.0 | -0.85 | 0.0 | 0.0 | 0.0 | 0.0 | 0.0 |
| 108 | R | 0.0 | 0.0 | 0.0 | 0.0 | 0.0 | 0.0 | 0.0 | 0.0 | 0.0 | 0.0 | 0.0 | 0.0 | 0.0 | 0.0 | 0.0 | 0.0 | 0.0 | 0.0 | 0.0 | 0.0 |
| 109 | A | 0.0 | -3.77 | 0.0 | 0.0 | -4.27 | -1.6 | -4.7 | -4.64 | -7.09 | -3.27 | -4.9 | -2.37 | -3.71 | -3.44 | -7.37 | -3.06 | -2.71 | -3.26 | 0.0 | -3.69 |
| 110 | A | 0.0 | -3.76 | 0.0 | 0.0 | -4.19 | -1.43 | -4.69 | -4.56 | -5.11 | -3.26 | -4.81 | -2.37 | -3.71 | -3.43 | -5.39 | -3.05 | -2.71 | -3.26 | 0.0 | -3.51 |
| 111 | E | -3.83 | -5.58 | -0.14 | 0.0 | -5.7 | -4.68 | -6.13 | -6.03 | -6.47 | -5.42 | -6.21 | -5.12 | -5.56 | -5.46 | -6.65 | -5.34 | -5.24 | -5.42 | -3.01 | -5.4 |
| 112 | D | -3.29 | -5.4 | 0.0 | 0.0 | -5.5 | -4.42 | -5.94 | -5.82 | -6.32 | -5.24 | -6.0 | -4.92 | -5.38 | -5.28 | -6.49 | -5.16 | -5.06 | -5.23 | -2.58 | -5.22 |
| 113 | D | -3.34 | -5.41 | 0.0 | 0.0 | -5.51 | -4.43 | -5.96 | -5.83 | -6.34 | -5.25 | -6.0 | -4.94 | -5.39 | -5.29 | -6.51 | -5.17 | -5.07 | -5.24 | -2.61 | -5.23 |
| 114 | E | -3.93 | -5.61 | -0.14 | 0.0 | -5.7 | -4.68 | -6.18 | -6.03 | -6.52 | -5.45 | -6.21 | -5.16 | -5.59 | -5.49 | -6.68 | -5.37 | -5.27 | -5.45 | -3.08 | -5.4 |
| 115 | D | -3.47 | -5.45 | 0.0 | 0.0 | -5.53 | -4.47 | -6.03 | -5.86 | -6.42 | -5.29 | -6.03 | -5.0 | -5.43 | -5.33 | -6.57 | -5.21 | -5.11 | -5.28 | -2.72 | -5.25 |
| 116 | D | -3.56 | -5.48 | 0.0 | 0.0 | -5.56 | -4.5 | -6.07 | -5.88 | -6.47 | -5.32 | -6.05 | -5.03 | -5.46 | -5.36 | -6.61 | -5.24 | -5.14 | -5.31 | -2.79 | -5.28 |
| 117 | D | -3.69 | -5.52 | 0.0 | 0.0 | -5.59 | -4.54 | -6.14 | -5.92 | -6.54 | -5.36 | -6.09 | -5.07 | -5.5 | -5.4 | -6.68 | -5.28 | -5.18 | -5.35 | -2.9 | -5.31 |
| 118 | V | 0.0 | -0.07 | -4.45 | -4.22 | -0.38 | 0.0 | -2.06 | -1.25 | -3.41 | 0.0 | -1.8 | 0.0 | -0.04 | 0.0 | -4.01 | 0.0 | 0.0 | 0.0 | 0.0 | 0.0 |
| 119 | D | -3.96 | -5.6 | 0.0 | 0.0 | -5.66 | -4.63 | -6.27 | -6.0 | -6.68 | -5.44 | -6.16 | -5.16 | -5.58 | -5.48 | -6.81 | -5.36 | -5.26 | -5.44 | -3.12 | -5.38 |
| 120 | T | 0.0 | -0.59 | 0.0 | 0.0 | -0.86 | 0.0 | -2.7 | -1.76 | -6.48 | -0.13 | -2.26 | 0.0 | -0.55 | -0.26 | -6.63 | 0.0 | 0.0 | -0.1 | 0.0 | -0.1 |
| 121 | K | 0.0 | 0.0 | 0.0 | 0.0 | 0.0 | 0.0 | 0.0 | 0.0 | 0.0 | 0.0 | 0.0 | 0.0 | 0.0 | 0.0 | -0.23 | 0.0 | 0.0 | 0.0 | 0.0 | 0.0 |
| 122 | K | 0.0 | 0.0 | 0.0 | 0.0 | 0.0 | 0.0 | 0.0 | 0.0 | 0.0 | 0.0 | 0.0 | 0.0 | 0.0 | 0.0 | -0.2 | 0.0 | 0.0 | 0.0 | 0.0 | 0.0 |
| 123 | Q | 0.0 | 0.0 | 0.0 | 0.0 | -0.08 | 0.0 | -2.18 | -1.0 | -7.85 | 0.0 | -1.44 | 0.0 | 0.0 | 0.0 | -7.91 | 0.0 | 0.0 | 0.0 | 0.0 | 0.0 |
| 124 | K | 0.0 | 0.0 | 0.0 | 0.0 | 0.0 | 0.0 | 0.0 | 0.0 | 0.0 | 0.0 | 0.0 | 0.0 | 0.0 | 0.0 | -0.12 | 0.0 | 0.0 | 0.0 | 0.0 | 0.0 |
| 125 | T | 0.0 | -0.55 | 0.0 | 0.0 | -0.68 | 0.0 | -2.85 | -1.62 | -4.37 | -0.11 | -2.04 | 0.0 | -0.48 | -0.21 | -4.53 | 0.0 | 0.0 | -0.05 | 0.0 | 0.0 |
| 126 | D | -4.28 | -5.73 | 0.0 | 0.0 | -5.72 | -4.72 | -6.46 | -6.09 | -6.97 | -5.58 | -6.2 | -5.32 | -5.7 | -5.6 | -7.02 | -5.47 | -5.41 | -5.56 | -3.36 | -5.42 |
| 127 | E | -4.53 | -5.92 | -0.15 | 0.0 | -5.88 | -4.92 | -6.6 | -6.26 | -7.09 | -5.76 | -6.35 | -5.51 | -5.88 | -5.77 | -7.13 | -5.64 | -5.59 | -5.74 | -3.73 | -5.55 |
| 128 | D | -4.32 | -5.72 | 0.0 | 0.0 | -5.65 | -4.64 | -6.45 | -6.04 | -7.0 | -5.58 | -6.11 | -5.34 | -5.68 | -5.58 | -7.01 | -5.45 | -5.42 | -5.54 | -3.23 | -5.34 |
| 129 | D | -4.38 | -5.76 | 0.0 | 0.0 | -5.7 | -4.69 | -6.51 | -6.08 | -7.05 | -5.62 | -6.16 | -5.38 | -5.72 | -5.62 | -7.06 | -5.49 | -5.46 | -5.59 | -3.36 | -5.38 |
