## Supplementary material for "Accelerated Missense Mutation Identification in Intrinsically Disordered Proteins using Deep Learning": Mutation heat-map Synuclein

| ID | WT | A | C | D | E | F | G | H | I | K | L | M | N | P | Q | R | S | T | V | W | Y |
|---|---|---|---|---|---|---|---|---|---|---|---|---|---|---|---|---|---|---|---|---|---|
| 1 | M | 0.1 | 0.01 | 0.12 | 0.14 | 0.02 | 0.05 | 0.04 | -0.01 | 0.1 | 0.05 | 0.0 | 0.05 | 0.04 | 0.06 | 0.03 | 0.04 | 0.06 | 0.06 | 0.09 | 0.06 |
| 2 | D | -0.8 | -1.09 | 0.0 | 0.02 | -1.47 | -1.34 | -2.0 | -1.44 | -2.51 | -0.98 | -1.55 | -0.91 | -1.03 | -1.0 | -2.96 | -1.03 | -0.92 | -0.96 | -1.33 | -1.44 |
| 3 | V | 0.08 | -0.05 | 0.5 | 0.51 | -0.23 | -0.16 | -0.4 | -0.22 | -0.64 | -0.0 | -0.27 | 0.02 | -0.02 | -0.01 | -1.1 | -0.02 | 0.02 | 0.0 | -0.15 | -0.21 |
| 4 | F | 0.25 | 0.13 | 0.62 | 0.62 | 0.0 | 0.05 | -0.13 | 0.0 | -0.33 | 0.18 | -0.02 | 0.2 | 0.16 | 0.17 | -0.73 | 0.16 | 0.2 | 0.18 | 0.09 | 0.03 |
| 5 | M | 0.29 | 0.17 | 0.64 | 0.65 | 0.04 | 0.08 | -0.09 | 0.04 | -0.28 | 0.22 | 0.0 | 0.24 | 0.2 | 0.21 | -0.7 | 0.2 | 0.24 | 0.22 | 0.12 | 0.06 |
| 6 | K | 0.45 | 0.36 | 0.75 | 0.76 | 0.22 | 0.25 | 0.15 | 0.22 | 0.0 | 0.39 | 0.18 | 0.41 | 0.37 | 0.38 | -0.39 | 0.37 | 0.4 | 0.39 | 0.28 | 0.23 |
| 7 | G | 0.18 | 0.04 | 0.57 | 0.58 | -0.04 | 0.0 | -0.27 | -0.05 | -0.5 | 0.08 | -0.08 | 0.12 | 0.06 | 0.07 | -0.83 | 0.06 | 0.11 | 0.09 | 0.04 | -0.02 |
| 8 | L | 0.1 | -0.04 | 0.52 | 0.53 | -0.22 | -0.15 | -0.4 | -0.2 | -0.63 | 0.0 | -0.27 | 0.03 | -0.02 | -0.01 | -1.11 | -0.02 | 0.02 | 0.01 | -0.14 | -0.21 |
| 9 | S | 0.13 | -0.02 | 0.54 | 0.55 | -0.11 | -0.06 | -0.36 | -0.11 | -0.59 | 0.03 | -0.15 | 0.06 | 0.01 | 0.02 | -0.95 | 0.0 | 0.05 | 0.03 | -0.03 | -0.1 |
| 10 | K | 0.45 | 0.35 | 0.75 | 0.75 | 0.29 | 0.32 | 0.14 | 0.29 | 0.0 | 0.38 | 0.27 | 0.41 | 0.36 | 0.38 | -0.22 | 0.36 | 0.4 | 0.38 | 0.34 | 0.3 |
| 11 | A | 0.0 | -0.16 | 0.46 | 0.47 | -0.29 | -0.22 | -0.58 | -0.27 | -0.95 | -0.11 | -0.34 | -0.07 | -0.14 | -0.12 | -1.47 | -0.14 | -0.08 | -0.11 | -0.2 | -0.28 |
| 12 | K | 0.44 | 0.34 | 0.74 | 0.74 | 0.27 | 0.3 | 0.14 | 0.27 | 0.0 | 0.38 | 0.24 | 0.4 | 0.36 | 0.36 | -0.25 | 0.35 | 0.39 | 0.38 | 0.33 | 0.28 |
| 13 | E | -0.83 | -1.19 | -0.02 | 0.0 | -1.63 | -1.48 | -2.13 | -1.57 | -2.55 | -1.06 | -1.74 | -0.97 | -1.13 | -1.11 | -3.03 | -1.14 | -1.0 | -1.06 | -1.46 | -1.62 |
| 14 | G | 0.16 | 0.01 | 0.55 | 0.56 | -0.05 | 0.0 | -0.29 | -0.04 | -0.48 | 0.06 | -0.09 | 0.09 | 0.03 | 0.04 | -0.78 | 0.03 | 0.08 | 0.06 | 0.04 | -0.03 |
| 15 | V | 0.11 | -0.05 | 0.51 | 0.52 | -0.24 | -0.17 | -0.4 | -0.22 | -0.6 | 0.0 | -0.29 | 0.04 | -0.02 | -0.01 | -1.08 | -0.03 | 0.02 | 0.0 | -0.15 | -0.23 |
| 16 | V | 0.11 | -0.05 | 0.51 | 0.52 | -0.16 | -0.09 | -0.4 | -0.14 | -0.59 | 0.0 | -0.21 | 0.04 | -0.02 | -0.01 | -0.96 | -0.03 | 0.02 | 0.0 | -0.08 | -0.15 |
| 17 | A | 0.0 | -0.17 | 0.44 | 0.45 | -0.29 | -0.22 | -0.56 | -0.27 | -0.77 | -0.11 | -0.34 | -0.08 | -0.14 | -0.13 | -1.2 | -0.14 | -0.09 | -0.12 | -0.2 | -0.28 |
| 18 | A | 0.0 | -0.17 | 0.44 | 0.45 | -0.29 | -0.21 | -0.56 | -0.26 | -0.76 | -0.11 | -0.34 | -0.08 | -0.14 | -0.13 | -1.18 | -0.15 | -0.09 | -0.12 | -0.19 | -0.27 |
| 19 | A | 0.0 | -0.17 | 0.43 | 0.44 | -0.28 | -0.21 | -0.56 | -0.26 | -0.75 | -0.11 | -0.33 | -0.08 | -0.14 | -0.13 | -1.16 | -0.15 | -0.09 | -0.12 | -0.19 | -0.27 |
| 20 | E | -0.8 | -1.15 | -0.02 | 0.0 | -1.4 | -1.25 | -2.06 | -1.33 | -2.48 | -1.02 | -1.5 | -0.93 | -1.1 | -1.07 | -2.84 | -1.11 | -0.96 | -1.03 | -1.24 | -1.4 |
| 21 | K | 0.43 | 0.32 | 0.72 | 0.72 | 0.27 | 0.3 | 0.12 | 0.27 | 0.0 | 0.36 | 0.24 | 0.38 | 0.34 | 0.34 | -0.22 | 0.33 | 0.37 | 0.36 | 0.32 | 0.27 |
| 22 | T | 0.09 | -0.07 | 0.5 | 0.5 | -0.18 | -0.11 | -0.44 | -0.15 | -0.75 | -0.02 | -0.23 | 0.01 | -0.05 | -0.04 | -1.2 | -0.05 | 0.0 | -0.02 | -0.09 | -0.17 |
| 23 | K | 0.43 | 0.32 | 0.72 | 0.73 | 0.25 | 0.28 | 0.12 | 0.25 | 0.0 | 0.36 | 0.22 | 0.38 | 0.34 | 0.35 | -0.26 | 0.33 | 0.38 | 0.36 | 0.31 | 0.26 |
| 24 | Q | 0.13 | -0.03 | 0.52 | 0.52 | -0.22 | -0.15 | -0.36 | -0.2 | -0.54 | 0.02 | -0.27 | 0.05 | -0.01 | 0.0 | -1.01 | -0.01 | 0.04 | 0.02 | -0.13 | -0.21 |
| 25 | G | 0.18 | 0.01 | 0.54 | 0.55 | -0.05 | 0.0 | -0.27 | -0.04 | -0.43 | 0.06 | -0.09 | 0.1 | 0.04 | 0.05 | -0.74 | 0.03 | 0.09 | 0.06 | 0.04 | -0.03 |
| 26 | V | 0.12 | -0.05 | 0.5 | 0.5 | -0.25 | -0.18 | -0.38 | -0.23 | -0.54 | 0.0 | -0.3 | 0.04 | -0.02 | -0.01 | -1.03 | -0.03 | 0.03 | 0.0 | -0.16 | -0.24 |
| 27 | A | 0.0 | -0.17 | 0.42 | 0.43 | -0.35 | -0.27 | -0.55 | -0.32 | -0.71 | -0.12 | -0.4 | -0.08 | -0.15 | -0.13 | -1.18 | -0.15 | -0.09 | -0.12 | -0.25 | -0.34 |
| 28 | E | -0.75 | -1.07 | -0.02 | 0.0 | -1.38 | -1.23 | -1.91 | -1.3 | -2.29 | -0.95 | -1.48 | -0.88 | -1.02 | -1.0 | -2.76 | -1.03 | -0.9 | -0.95 | -1.22 | -1.38 |
| 29 | A | 0.0 | -0.18 | 0.41 | 0.42 | -0.35 | -0.27 | -0.55 | -0.32 | -0.7 | -0.12 | -0.4 | -0.08 | -0.15 | -0.13 | -1.16 | -0.15 | -0.09 | -0.12 | -0.25 | -0.34 |
| 30 | A | 0.0 | -0.18 | 0.42 | 0.43 | -0.43 | -0.35 | -0.55 | -0.41 | -0.71 | -0.12 | -0.49 | -0.08 | -0.15 | -0.13 | -1.3 | -0.15 | -0.09 | -0.12 | -0.32 | -0.42 |
| 31 | G | 0.19 | 0.02 | 0.54 | 0.55 | -0.05 | 0.0 | -0.24 | -0.04 | -0.39 | 0.08 | -0.1 | 0.11 | 0.05 | 0.06 | -0.72 | 0.04 | 0.1 | 0.07 | 0.04 | -0.04 |
| 32 | K | 0.42 | 0.3 | 0.7 | 0.7 | 0.18 | 0.22 | 0.1 | 0.18 | 0.0 | 0.34 | 0.14 | 0.36 | 0.32 | 0.32 | -0.36 | 0.31 | 0.36 | 0.34 | 0.25 | 0.19 |
| 33 | T | 0.09 | -0.08 | 0.48 | 0.49 | -0.23 | -0.16 | -0.42 | -0.2 | -0.69 | -0.02 | -0.28 | 0.02 | -0.05 | -0.04 | -1.22 | -0.06 | 0.0 | -0.02 | -0.14 | -0.22 |
| 34 | K | 0.41 | 0.29 | 0.68 | 0.69 | 0.2 | 0.24 | 0.1 | 0.21 | 0.0 | 0.33 | 0.17 | 0.36 | 0.31 | 0.32 | -0.29 | 0.3 | 0.35 | 0.33 | 0.27 | 0.21 |
| 35 | E | -0.72 | -1.05 | -0.02 | 0.0 | -1.5 | -1.34 | -1.85 | -1.42 | -2.2 | -0.92 | -1.61 | -0.86 | -0.99 | -0.97 | -2.83 | -1.01 | -0.88 | -0.93 | -1.32 | -1.5 |
| 36 | G | 0.2 | 0.04 | 0.53 | 0.55 | -0.05 | 0.0 | -0.2 | -0.04 | -0.32 | 0.09 | -0.1 | 0.13 | 0.06 | 0.08 | -0.67 | 0.06 | 0.12 | 0.09 | 0.04 | -0.04 |
| 37 | V | 0.12 | -0.05 | 0.48 | 0.49 | -0.28 | -0.21 | -0.36 | -0.25 | -0.48 | 0.01 | -0.34 | 0.04 | -0.02 | -0.01 | -0.99 | -0.03 | 0.03 | 0.0 | -0.18 | -0.27 |
| 38 | L | 0.12 | -0.05 | 0.48 | 0.49 | -0.28 | -0.2 | -0.35 | -0.25 | -0.48 | 0.0 | -0.34 | 0.04 | -0.03 | -0.01 | -0.99 | -0.03 | 0.03 | 0.0 | -0.18 | -0.27 |
| 39 | Y | 0.26 | 0.1 | 0.58 | 0.58 | -0.01 | 0.04 | -0.13 | -0.0 | -0.23 | 0.16 | -0.05 | 0.19 | 0.13 | 0.14 | -0.59 | 0.12 | 0.18 | 0.15 | 0.08 | 0.0 |
| 40 | V | 0.12 | -0.05 | 0.47 | 0.48 | -0.34 | -0.26 | -0.35 | -0.31 | -0.47 | 0.01 | -0.4 | 0.04 | -0.02 | -0.01 | -1.06 | -0.03 | 0.03 | 0.0 | -0.24 | -0.33 |
| 41 | G | 0.22 | 0.06 | 0.54 | 0.55 | -0.05 | 0.0 | -0.18 | -0.04 | -0.28 | 0.11 | -0.1 | 0.14 | 0.08 | 0.09 | -0.66 | 0.07 | 0.13 | 0.1 | 0.04 | -0.04 |
| 42 | S | 0.15 | -0.01 | 0.5 | 0.5 | -0.21 | -0.13 | -0.29 | -0.18 | -0.41 | 0.04 | -0.26 | 0.08 | 0.01 | 0.02 | -0.89 | 0.0 | 0.06 | 0.03 | -0.12 | -0.2 |
| 43 | K | 0.39 | 0.27 | 0.66 | 0.67 | 0.18 | 0.22 | 0.08 | 0.19 | 0.0 | 0.31 | 0.14 | 0.34 | 0.29 | 0.3 | -0.29 | 0.29 | 0.33 | 0.31 | 0.25 | 0.2 |
| 44 | T | 0.09 | -0.08 | 0.46 | 0.46 | -0.22 | -0.14 | -0.39 | -0.18 | -0.63 | -0.02 | -0.27 | 0.02 | -0.05 | -0.04 | -1.09 | -0.06 | 0.0 | -0.02 | -0.12 | -0.21 |
| 45 | K | 0.38 | 0.26 | 0.65 | 0.65 | 0.17 | 0.21 | 0.08 | 0.18 | 0.0 | 0.3 | 0.13 | 0.33 | 0.28 | 0.29 | -0.29 | 0.27 | 0.32 | 0.3 | 0.24 | 0.18 |
| 46 | E | -0.65 | -0.95 | -0.02 | 0.0 | -1.37 | -1.21 | -1.64 | -1.29 | -1.9 | -0.85 | -1.48 | -0.78 | -0.91 | -0.88 | -2.67 | -0.92 | -0.81 | -0.86 | -1.2 | -1.37 |
| 47 | G | 0.2 | 0.03 | 0.51 | 0.52 | -0.05 | 0.0 | -0.18 | -0.04 | -0.26 | 0.08 | -0.1 | 0.12 | 0.05 | 0.07 | -0.6 | 0.05 | 0.11 | 0.08 | 0.04 | -0.04 |
| 48 | V | 0.12 | -0.05 | 0.45 | 0.46 | -0.27 | -0.2 | -0.32 | -0.24 | -0.41 | 0.01 | -0.33 | 0.04 | -0.02 | -0.01 | -0.9 | -0.03 | 0.03 | 0.0 | -0.18 | -0.27 |
| 49 | V | 0.12 | -0.05 | 0.45 | 0.46 | -0.22 | -0.14 | -0.3 | -0.18 | -0.41 | 0.01 | -0.27 | 0.04 | -0.02 | -0.01 | -0.83 | -0.03 | 0.03 | 0.0 | -0.12 | -0.21 |
| 50 | H | 0.34 | 0.21 | 0.61 | 0.62 | 0.05 | 0.1 | 0.0 | 0.06 | -0.04 | 0.25 | 0.01 | 0.29 | 0.23 | 0.24 | -0.43 | 0.23 | 0.27 | 0.25 | 0.14 | 0.06 |
| 51 | G | 0.2 | 0.03 | 0.5 | 0.51 | -0.05 | 0.0 | -0.18 | -0.04 | -0.25 | 0.08 | -0.1 | 0.12 | 0.05 | 0.06 | -0.58 | 0.04 | 0.1 | 0.08 | 0.04 | -0.04 |
| 52 | V | 0.12 | -0.05 | 0.44 | 0.45 | -0.26 | -0.18 | -0.3 | -0.23 | -0.38 | 0.01 | -0.31 | 0.04 | -0.02 | -0.01 | -0.85 | -0.03 | 0.03 | 0.0 | -0.16 | -0.25 |
| 53 | A | 0.0 | -0.18 | 0.35 | 0.36 | -0.36 | -0.28 | -0.49 | -0.32 | -0.56 | -0.12 | -0.42 | -0.08 | -0.16 | -0.14 | -0.95 | -0.16 | -0.1 | -0.13 | -0.25 | -0.35 |
| 54 | T | 0.1 | -0.08 | 0.41 | 0.42 | -0.19 | -0.12 | -0.34 | -0.16 | -0.4 | -0.02 | -0.25 | 0.02 | -0.06 | -0.04 | -0.75 | -0.06 | 0.0 | -0.03 | -0.1 | -0.19 |
| 55 | V | 0.12 | -0.05 | 0.43 | 0.44 | -0.14 | -0.08 | -0.29 | -0.12 | -0.35 | 0.01 | -0.19 | 0.04 | -0.02 | -0.01 | -0.68 | -0.03 | 0.03 | 0.0 | -0.06 | -0.14 |
| 56 | A | 0.0 | -0.18 | 0.34 | 0.35 | -0.31 | -0.23 | -0.47 | -0.28 | -0.52 | -0.12 | -0.37 | -0.08 | -0.16 | -0.14 | -0.86 | -0.16 | -0.1 | -0.13 | -0.21 | -0.3 |
| 57 | E | -0.55 | -0.84 | -0.02 | 0.0 | -0.99 | -0.89 | -1.35 | -0.96 | -1.49 | -0.75 | -1.08 | -0.68 | -0.81 | -0.79 | -2.15 | -0.81 | -0.71 | -0.76 | -0.86 | -0.98 |
| 58 | K | 0.33 | 0.21 | 0.57 | 0.58 | 0.14 | 0.19 | 0.04 | 0.15 | 0.0 | 0.25 | 0.1 | 0.28 | 0.22 | 0.24 | -0.23 | 0.22 | 0.27 | 0.25 | 0.22 | 0.15 |
| 59 | T | 0.1 | -0.08 | 0.4 | 0.42 | -0.17 | -0.1 | -0.33 | -0.14 | -0.5 | -0.02 | -0.22 | 0.02 | -0.06 | -0.04 | -0.82 | -0.06 | 0.0 | -0.03 | -0.08 | -0.17 |
| 60 | K | 0.33 | 0.2 | 0.56 | 0.57 | 0.13 | 0.18 | 0.03 | 0.14 | 0.0 | 0.24 | 0.09 | 0.27 | 0.22 | 0.23 | -0.23 | 0.21 | 0.26 | 0.24 | 0.21 | 0.14 |
| 61 | E | -0.55 | -0.84 | -0.02 | 0.0 | -0.98 | -0.89 | -1.24 | -0.96 | -1.34 | -0.75 | -1.06 | -0.68 | -0.8 | -0.78 | -1.99 | -0.81 | -0.7 | -0.75 | -0.83 | -0.96 |
| 62 | Q | 0.14 | -0.03 | 0.42 | 0.43 | -0.11 | -0.06 | -0.21 | -0.1 | -0.23 | 0.02 | -0.16 | 0.06 | -0.01 | 0.0 | -0.55 | -0.01 | 0.04 | 0.02 | -0.02 | -0.11 |
| 63 | V | 0.12 | -0.05 | 0.42 | 0.43 | -0.14 | -0.08 | -0.22 | -0.12 | -0.24 | 0.01 | -0.2 | 0.04 | -0.02 | -0.01 | -0.56 | -0.03 | 0.03 | 0.0 | -0.06 | -0.14 |
| 64 | T | 0.1 | -0.08 | 0.4 | 0.42 | -0.2 | -0.12 | -0.24 | -0.16 | -0.25 | -0.02 | -0.25 | 0.02 | -0.06 | -0.04 | -0.59 | -0.06 | 0.0 | -0.03 | -0.11 | -0.19 |
| 65 | N | 0.08 | -0.09 | 0.4 | 0.41 | -0.24 | -0.16 | -0.23 | -0.21 | -0.25 | -0.03 | -0.29 | 0.0 | -0.07 | -0.05 | -0.61 | -0.07 | -0.01 | -0.04 | -0.14 | -0.24 |
| 66 | V | 0.12 | -0.05 | 0.43 | 0.44 | -0.26 | -0.18 | -0.16 | -0.23 | -0.16 | 0.01 | -0.32 | 0.04 | -0.02 | -0.01 | -0.6 | -0.03 | 0.03 | 0.0 | -0.16 | -0.26 |
| 67 | G | 0.26 | 0.1 | 0.53 | 0.53 | -0.05 | 0.0 | 0.02 | -0.04 | -0.0 | 0.16 | -0.1 | 0.19 | 0.13 | 0.14 | -0.32 | 0.12 | 0.18 | 0.15 | 0.04 | -0.04 |
| 68 | G | 0.26 | 0.11 | 0.53 | 0.53 | -0.05 | 0.0 | 0.02 | -0.04 | 0.01 | 0.16 | -0.1 | 0.2 | 0.13 | 0.14 | -0.31 | 0.12 | 0.18 | 0.16 | 0.04 | -0.04 |
| 69 | A | 0.0 | -0.18 | 0.35 | 0.36 | -0.46 | -0.38 | -0.3 | -0.43 | -0.3 | -0.12 | -0.52 | -0.08 | -0.16 | -0.14 | -0.79 | -0.16 | -0.1 | -0.13 | -0.35 | -0.45 |
| 70 | V | 0.12 | -0.05 | 0.42 | 0.44 | -0.2 | -0.13 | -0.12 | -0.17 | -0.13 | 0.01 | -0.25 | 0.04 | -0.02 | -0.01 | -0.49 | -0.03 | 0.03 | 0.0 | -0.11 | -0.2 |
| 71 | V | 0.12 | -0.05 | 0.42 | 0.43 | -0.2 | -0.13 | -0.12 | -0.17 | -0.12 | 0.01 | -0.25 | 0.04 | -0.02 | -0.01 | -0.48 | -0.03 | 0.03 | 0.0 | -0.11 | -0.2 |
| 72 | T | 0.1 | -0.08 | 0.4 | 0.41 | -0.31 | -0.24 | -0.16 | -0.28 | -0.15 | -0.02 | -0.37 | 0.02 | -0.06 | -0.04 | -0.58 | -0.06 | 0.0 | -0.03 | -0.22 | -0.31 |
| 73 | G | 0.2 | 0.02 | 0.47 | 0.48 | -0.05 | 0.0 | -0.04 | -0.04 | -0.04 | 0.08 | -0.1 | 0.12 | 0.05 | 0.06 | -0.27 | 0.04 | 0.1 | 0.08 | 0.04 | -0.04 |
| 74 | V | 0.12 | -0.05 | 0.41 | 0.42 | -0.26 | -0.19 | -0.11 | -0.23 | -0.11 | 0.01 | -0.32 | 0.04 | -0.02 | -0.01 | -0.52 | -0.03 | 0.03 | 0.0 | -0.17 | -0.26 |
| 75 | T | 0.1 | -0.08 | 0.38 | 0.39 | -0.23 | -0.15 | -0.16 | -0.19 | -0.14 | -0.02 | -0.28 | 0.02 | -0.06 | -0.04 | -0.49 | -0.06 | 0.0 | -0.03 | -0.13 | -0.22 |
| 76 | A | 0.0 | -0.18 | 0.31 | 0.32 | -0.34 | -0.26 | -0.3 | -0.31 | -0.27 | -0.12 | -0.4 | -0.08 | -0.16 | -0.14 | -0.62 | -0.16 | -0.1 | -0.13 | -0.24 | -0.33 |
| 77 | V | 0.12 | -0.05 | 0.38 | 0.39 | -0.15 | -0.08 | -0.15 | -0.13 | -0.12 | 0.01 | -0.2 | 0.04 | -0.02 | -0.01 | -0.43 | -0.03 | 0.03 | 0.0 | -0.06 | -0.15 |
| 78 | A | 0.0 | -0.18 | 0.28 | 0.29 | -0.32 | -0.25 | -0.34 | -0.29 | -0.3 | -0.12 | -0.38 | -0.08 | -0.16 | -0.14 | -0.63 | -0.16 | -0.1 | -0.13 | -0.22 | -0.32 |
| 79 | Q | 0.14 | -0.03 | 0.37 | 0.38 | -0.12 | -0.06 | -0.15 | -0.11 | -0.13 | 0.02 | -0.17 | 0.06 | -0.01 | 0.0 | -0.43 | -0.01 | 0.04 | 0.02 | -0.03 | -0.12 |
| 80 | K | 0.21 | 0.04 | 0.44 | 0.45 | -0.04 | 0.01 | -0.02 | -0.04 | 0.0 | 0.1 | -0.09 | 0.14 | 0.06 | 0.08 | -0.25 | 0.06 | 0.12 | 0.09 | 0.06 | -0.03 |
| 81 | T | 0.1 | -0.08 | 0.32 | 0.33 | -0.2 | -0.13 | -0.24 | -0.17 | -0.18 | -0.02 | -0.26 | 0.02 | -0.06 | -0.04 | -0.53 | -0.06 | 0.0 | -0.03 | -0.11 | -0.2 |
| 82 | V | 0.12 | -0.05 | 0.32 | 0.34 | -0.18 | -0.11 | -0.18 | -0.15 | -0.13 | 0.01 | -0.24 | 0.04 | -0.02 | -0.01 | -0.48 | -0.03 | 0.03 | 0.0 | -0.1 | -0.18 |
| 83 | E | -0.31 | -0.58 | -0.02 | 0.0 | -0.88 | -0.79 | -0.78 | -0.86 | -0.7 | -0.5 | -0.96 | -0.43 | -0.55 | -0.53 | -1.31 | -0.56 | -0.46 | -0.5 | -0.74 | -0.87 |
| 84 | G | 0.22 | 0.06 | 0.39 | 0.4 | -0.05 | 0.0 | -0.05 | -0.04 | -0.01 | 0.11 | -0.1 | 0.15 | 0.08 | 0.09 | -0.27 | 0.07 | 0.13 | 0.1 | 0.04 | -0.04 |
| 85 | A | 0.0 | -0.18 | 0.22 | 0.24 | -0.55 | -0.46 | -0.35 | -0.52 | -0.27 | -0.12 | -0.61 | -0.08 | -0.16 | -0.14 | -0.85 | -0.16 | -0.1 | -0.13 | -0.43 | -0.54 |
| 86 | G | 0.24 | 0.08 | 0.4 | 0.41 | -0.05 | 0.0 | -0.02 | -0.04 | 0.02 | 0.13 | -0.1 | 0.17 | 0.1 | 0.11 | -0.27 | 0.1 | 0.16 | 0.13 | 0.04 | -0.04 |
| 87 | S | 0.16 | -0.01 | 0.33 | 0.34 | -0.3 | -0.21 | -0.11 | -0.26 | -0.07 | 0.04 | -0.35 | 0.08 | 0.01 | 0.02 | -0.56 | 0.0 | 0.06 | 0.03 | -0.2 | -0.29 |
| 88 | I | 0.26 | 0.11 | 0.41 | 0.42 | -0.0 | 0.05 | 0.01 | 0.0 | 0.06 | 0.16 | -0.04 | 0.2 | 0.13 | 0.14 | -0.2 | 0.12 | 0.18 | 0.15 | 0.09 | 0.01 |
| 89 | A | 0.0 | -0.18 | 0.21 | 0.22 | -0.46 | -0.38 | -0.33 | -0.43 | -0.24 | -0.12 | -0.53 | -0.08 | -0.16 | -0.14 | -0.72 | -0.16 | -0.1 | -0.13 | -0.35 | -0.46 |
| 90 | A | 0.0 | -0.18 | 0.2 | 0.22 | -0.39 | -0.3 | -0.33 | -0.36 | -0.23 | -0.12 | -0.45 | -0.08 | -0.16 | -0.14 | -0.63 | -0.16 | -0.1 | -0.13 | -0.28 | -0.38 |
| 91 | A | 0.0 | -0.18 | 0.2 | 0.21 | -0.39 | -0.3 | -0.32 | -0.36 | -0.22 | -0.12 | -0.45 | -0.08 | -0.16 | -0.14 | -0.62 | -0.16 | -0.1 | -0.13 | -0.28 | -0.38 |
| 92 | T | 0.1 | -0.08 | 0.27 | 0.28 | -0.32 | -0.24 | -0.18 | -0.28 | -0.1 | -0.02 | -0.37 | 0.02 | -0.06 | -0.04 | -0.53 | -0.06 | 0.0 | -0.03 | -0.22 | -0.31 |
| 93 | G | 0.26 | 0.11 | 0.4 | 0.41 | -0.05 | 0.0 | 0.03 | -0.04 | 0.09 | 0.16 | -0.1 | 0.2 | 0.13 | 0.14 | -0.21 | 0.12 | 0.18 | 0.16 | 0.04 | -0.04 |
| 94 | F | 0.3 | 0.16 | 0.43 | 0.44 | 0.0 | 0.06 | 0.09 | 0.02 | 0.15 | 0.21 | -0.04 | 0.25 | 0.18 | 0.2 | -0.12 | 0.18 | 0.24 | 0.21 | 0.1 | 0.02 |
| 95 | V | 0.12 | -0.05 | 0.28 | 0.29 | -0.27 | -0.19 | -0.13 | -0.24 | -0.15 | 0.01 | -0.32 | 0.04 | -0.02 | -0.01 | -0.59 | -0.03 | 0.03 | 0.0 | -0.17 | -0.26 |
| 96 | K | 0.18 | 0.01 | 0.34 | 0.36 | -0.12 | -0.06 | -0.08 | -0.09 | 0.0 | 0.06 | -0.16 | 0.1 | 0.02 | 0.04 | -0.29 | 0.02 | 0.09 | 0.06 | -0.03 | -0.11 |
| 97 | K | 0.16 | -0.01 | 0.32 | 0.33 | -0.12 | -0.06 | -0.09 | -0.1 | 0.0 | 0.04 | -0.16 | 0.09 | 0.01 | 0.02 | -0.27 | 0.01 | 0.07 | 0.04 | -0.03 | -0.11 |
| 98 | D | -0.16 | -0.41 | 0.0 | 0.02 | -0.59 | -0.5 | -0.52 | -0.56 | -0.51 | -0.33 | -0.66 | -0.26 | -0.38 | -0.36 | -0.9 | -0.38 | -0.29 | -0.33 | -0.47 | -0.58 |
| 99 | Q | 0.14 | -0.03 | 0.25 | 0.27 | -0.16 | -0.09 | -0.08 | -0.14 | 0.0 | 0.02 | -0.22 | 0.06 | -0.01 | 0.0 | -0.29 | -0.01 | 0.04 | 0.02 | -0.07 | -0.16 |
| 100 | L | 0.12 | -0.05 | 0.24 | 0.25 | -0.27 | -0.19 | -0.1 | -0.24 | -0.01 | 0.0 | -0.32 | 0.04 | -0.03 | -0.01 | -0.39 | -0.03 | 0.03 | 0.0 | -0.17 | -0.26 |
| 101 | G | 0.2 | 0.03 | 0.3 | 0.31 | -0.05 | 0.0 | -0.02 | -0.04 | 0.07 | 0.08 | -0.1 | 0.12 | 0.05 | 0.06 | -0.13 | 0.04 | 0.1 | 0.08 | 0.04 | -0.04 |
| 102 | K | 0.12 | -0.06 | 0.22 | 0.24 | -0.28 | -0.2 | -0.1 | -0.25 | 0.0 | -0.01 | -0.34 | 0.04 | -0.04 | -0.02 | -0.37 | -0.04 | 0.02 | -0.01 | -0.18 | -0.28 |
| 103 | N | 0.08 | -0.09 | 0.28 | 0.3 | -0.26 | -0.18 | -0.13 | -0.23 | -0.02 | -0.03 | -0.32 | 0.0 | -0.07 | -0.05 | -0.33 | -0.07 | -0.01 | -0.04 | -0.16 | -0.25 |
| 104 | E | -0.1 | -0.32 | -0.02 | 0.0 | -0.54 | -0.45 | -0.38 | -0.51 | -0.19 | -0.24 | -0.6 | -0.19 | -0.29 | -0.27 | -0.58 | -0.3 | -0.2 | -0.25 | -0.42 | -0.53 |
| 105 | E | -0.09 | -0.3 | -0.02 | 0.0 | -0.6 | -0.51 | -0.35 | -0.57 | -0.16 | -0.23 | -0.66 | -0.18 | -0.27 | -0.26 | -0.62 | -0.28 | -0.19 | -0.23 | -0.48 | -0.59 |
| 106 | G | 0.2 | 0.03 | 0.35 | 0.36 | -0.05 | 0.0 | 0.01 | -0.04 | 0.14 | 0.08 | -0.1 | 0.12 | 0.05 | 0.06 | -0.06 | 0.05 | 0.11 | 0.08 | 0.04 | -0.04 |
| 107 | A | 0.0 | -0.18 | 0.06 | 0.08 | -0.46 | -0.38 | -0.22 | -0.43 | -0.06 | -0.12 | -0.52 | -0.08 | -0.16 | -0.14 | -0.48 | -0.16 | -0.1 | -0.13 | -0.35 | -0.45 |
| 108 | P | 0.15 | -0.02 | 0.2 | 0.22 | -0.16 | -0.08 | -0.03 | -0.13 | 0.1 | 0.03 | -0.21 | 0.07 | 0.0 | 0.02 | -0.16 | 0.0 | 0.06 | 0.03 | -0.07 | -0.16 |
| 109 | Q | 0.14 | -0.03 | 0.18 | 0.19 | -0.17 | -0.1 | -0.04 | -0.15 | 0.1 | 0.02 | -0.22 | 0.06 | -0.01 | 0.0 | -0.16 | -0.01 | 0.04 | 0.02 | -0.08 | -0.17 |
| 110 | E | -0.05 | -0.25 | -0.02 | 0.0 | -0.54 | -0.45 | -0.27 | -0.51 | -0.09 | -0.18 | -0.6 | -0.14 | -0.22 | -0.2 | -0.5 | -0.23 | -0.15 | -0.18 | -0.42 | -0.53 |
| 111 | G | 0.26 | 0.1 | 0.27 | 0.29 | -0.05 | 0.0 | 0.11 | -0.04 | 0.24 | 0.16 | -0.1 | 0.19 | 0.13 | 0.14 | -0.02 | 0.12 | 0.18 | 0.15 | 0.04 | -0.04 |
| 112 | I | 0.29 | 0.15 | 0.3 | 0.32 | -0.0 | 0.05 | 0.16 | 0.0 | 0.27 | 0.2 | -0.04 | 0.23 | 0.17 | 0.18 | 0.03 | 0.17 | 0.22 | 0.2 | 0.09 | 0.01 |
| 113 | L | 0.12 | -0.05 | 0.25 | 0.26 | -0.27 | -0.19 | -0.03 | -0.24 | 0.11 | 0.0 | -0.33 | 0.04 | -0.03 | -0.01 | -0.21 | -0.03 | 0.03 | 0.0 | -0.18 | -0.27 |
| 114 | E | -0.01 | -0.2 | -0.02 | 0.0 | -0.42 | -0.34 | -0.19 | -0.39 | -0.01 | -0.14 | -0.48 | -0.1 | -0.17 | -0.16 | -0.34 | -0.18 | -0.11 | -0.14 | -0.31 | -0.41 |
| 115 | D | 0.01 | -0.16 | 0.0 | 0.02 | -0.44 | -0.36 | -0.15 | -0.41 | 0.02 | -0.11 | -0.5 | -0.07 | -0.14 | -0.13 | -0.35 | -0.14 | -0.08 | -0.11 | -0.33 | -0.43 |
| 116 | M | 0.27 | 0.12 | 0.36 | 0.38 | 0.04 | 0.1 | 0.14 | 0.05 | 0.28 | 0.17 | 0.0 | 0.21 | 0.14 | 0.16 | 0.11 | 0.14 | 0.2 | 0.17 | 0.13 | 0.06 |
| 117 | P | 0.15 | -0.02 | 0.14 | 0.16 | -0.22 | -0.14 | 0.0 | -0.18 | 0.16 | 0.03 | -0.27 | 0.07 | 0.0 | 0.02 | -0.13 | 0.0 | 0.06 | 0.03 | -0.12 | -0.21 |
| 118 | V | 0.12 | -0.05 | 0.1 | 0.12 | -0.18 | -0.1 | -0.02 | -0.15 | 0.14 | 0.01 | -0.23 | 0.04 | -0.02 | -0.01 | -0.08 | -0.03 | 0.03 | 0.0 | -0.09 | -0.18 |
| 119 | D | 0.02 | -0.15 | 0.0 | 0.02 | -0.3 | -0.22 | -0.12 | -0.27 | 0.05 | -0.1 | -0.35 | -0.05 | -0.13 | -0.11 | -0.19 | -0.13 | -0.07 | -0.1 | -0.2 | -0.29 |
| 120 | P | 0.15 | -0.02 | 0.24 | 0.25 | -0.11 | -0.05 | 0.01 | -0.1 | 0.18 | 0.03 | -0.16 | 0.07 | 0.0 | 0.02 | -0.01 | 0.0 | 0.06 | 0.03 | -0.02 | -0.1 |
| 121 | D | 0.03 | -0.14 | 0.0 | 0.02 | -0.27 | -0.2 | -0.1 | -0.24 | 0.06 | -0.09 | -0.33 | -0.04 | -0.12 | -0.1 | -0.16 | -0.12 | -0.06 | -0.09 | -0.18 | -0.27 |
| 122 | N | 0.08 | -0.09 | 0.18 | 0.2 | -0.21 | -0.13 | -0.05 | -0.18 | 0.12 | -0.03 | -0.26 | 0.0 | -0.06 | -0.05 | -0.08 | -0.06 | -0.01 | -0.04 | -0.12 | -0.2 |
| 123 | E | 0.02 | -0.16 | -0.02 | 0.0 | -0.32 | -0.25 | -0.12 | -0.3 | 0.05 | -0.1 | -0.38 | -0.06 | -0.14 | -0.12 | -0.2 | -0.14 | -0.08 | -0.11 | -0.23 | -0.32 |
| 124 | A | 0.0 | -0.18 | -0.04 | -0.02 | -0.44 | -0.35 | -0.14 | -0.41 | 0.04 | -0.12 | -0.5 | -0.08 | -0.15 | -0.14 | -0.3 | -0.15 | -0.09 | -0.12 | -0.33 | -0.43 |
| 125 | Y | 0.25 | 0.09 | 0.22 | 0.24 | -0.01 | 0.04 | 0.12 | -0.0 | 0.27 | 0.14 | -0.05 | 0.18 | 0.11 | 0.13 | 0.08 | 0.11 | 0.17 | 0.14 | 0.08 | 0.0 |
| 126 | E | 0.02 | -0.16 | -0.02 | 0.0 | -0.5 | -0.41 | -0.12 | -0.47 | 0.05 | -0.1 | -0.56 | -0.06 | -0.13 | -0.12 | -0.36 | -0.14 | -0.08 | -0.1 | -0.38 | -0.48 |
| 127 | M | 0.28 | 0.14 | 0.25 | 0.27 | 0.04 | 0.09 | 0.18 | 0.05 | 0.31 | 0.19 | 0.0 | 0.22 | 0.16 | 0.18 | 0.13 | 0.16 | 0.22 | 0.19 | 0.13 | 0.06 |
| 128 | P | 0.14 | -0.02 | 0.11 | 0.13 | -0.22 | -0.14 | 0.01 | -0.19 | 0.17 | 0.03 | -0.27 | 0.07 | 0.0 | 0.02 | -0.1 | 0.0 | 0.06 | 0.03 | -0.12 | -0.21 |
| 129 | S | 0.14 | -0.02 | 0.23 | 0.25 | -0.14 | -0.08 | 0.02 | -0.13 | 0.18 | 0.03 | -0.19 | 0.07 | 0.01 | 0.02 | -0.03 | 0.0 | 0.06 | 0.03 | -0.06 | -0.14 |
| 130 | E | 0.02 | -0.15 | -0.02 | 0.0 | -0.32 | -0.24 | -0.1 | -0.29 | 0.06 | -0.09 | -0.37 | -0.05 | -0.12 | -0.11 | -0.19 | -0.12 | -0.07 | -0.09 | -0.22 | -0.31 |
| 131 | E | 0.02 | -0.15 | -0.02 | 0.0 | -0.39 | -0.31 | -0.1 | -0.37 | 0.06 | -0.09 | -0.45 | -0.06 | -0.12 | -0.11 | -0.26 | -0.12 | -0.07 | -0.09 | -0.29 | -0.38 |
| 132 | G | 0.24 | 0.1 | 0.32 | 0.34 | -0.05 | 0.0 | 0.13 | -0.05 | 0.27 | 0.14 | -0.09 | 0.18 | 0.12 | 0.13 | 0.04 | 0.12 | 0.17 | 0.14 | 0.04 | -0.04 |
| 133 | Y | 0.28 | 0.14 | 0.26 | 0.27 | -0.01 | 0.04 | 0.17 | -0.01 | 0.3 | 0.19 | -0.05 | 0.22 | 0.16 | 0.17 | 0.06 | 0.16 | 0.21 | 0.19 | 0.08 | 0.0 |
| 134 | Q | 0.12 | -0.03 | 0.1 | 0.12 | -0.24 | -0.16 | -0.0 | -0.22 | 0.15 | 0.02 | -0.29 | 0.05 | -0.01 | 0.0 | -0.13 | -0.01 | 0.04 | 0.02 | -0.15 | -0.23 |
| 135 | D | 0.03 | -0.14 | 0.0 | 0.02 | -0.39 | -0.31 | -0.1 | -0.37 | 0.05 | -0.08 | -0.44 | -0.05 | -0.11 | -0.1 | -0.29 | -0.11 | -0.06 | -0.08 | -0.29 | -0.38 |
| 136 | Y | 0.21 | 0.05 | 0.3 | 0.31 | -0.01 | 0.04 | 0.09 | -0.01 | 0.23 | 0.1 | -0.05 | 0.14 | 0.08 | 0.09 | 0.06 | 0.08 | 0.13 | 0.1 | 0.07 | 0.0 |
| 137 | E | -0.0 | -0.16 | -0.02 | 0.0 | -0.39 | -0.31 | -0.13 | -0.37 | 0.02 | -0.11 | -0.44 | -0.08 | -0.14 | -0.12 | -0.3 | -0.14 | -0.09 | -0.11 | -0.29 | -0.37 |
| 138 | P | 0.13 | -0.02 | 0.23 | 0.25 | -0.1 | -0.05 | 0.01 | -0.1 | 0.14 | 0.02 | -0.14 | 0.06 | 0.0 | 0.02 | -0.03 | 0.0 | 0.05 | 0.03 | -0.03 | -0.09 |
| 139 | E | -0.01 | -0.17 | -0.02 | 0.0 | -0.26 | -0.19 | -0.14 | -0.25 | -0.01 | -0.12 | -0.3 | -0.09 | -0.14 | -0.12 | -0.2 | -0.14 | -0.09 | -0.11 | -0.18 | -0.25 |
| 140 | A | 0.0 | -0.15 | 0.0 | 0.02 | -0.23 | -0.16 | -0.13 | -0.22 | 0.0 | -0.1 | -0.27 | -0.07 | -0.12 | -0.11 | -0.18 | -0.12 | -0.08 | -0.1 | -0.15 | -0.22 |
